# Pregnenolone reorganizes cytoskeleton to promote neuron development via CLIP1

**DOI:** 10.1101/2022.08.01.502406

**Authors:** Kolas Viktoryia, Yi-Ting Wu, Jose Sandino A. Bandonil, Bon-Chu Chung

## Abstract

Pregnenolone (P5) is a neurosteroid produced in the brain. It improves cognitive function and protects against cannabis intoxication as well as spinal cord injury. P5 activates CLIP1, which helps microtubule polymerization at its growing end; however, the significance of P5 activation of CLIP1 in the brain is still unknown. Here we examined the roles of P5 in cultured neurons and in zebrafish cerebellum. We show that P5 promotes neurite outgrowth and facilitates axon development of cultured cerebellar granule neurons. P5 also changes the morphology of axon growth cone and promotes dynamic microtubule invasion into the distal part of filopodia at the growth cone. We have used CRISPR to disrupt *clip1a* in zebrafish, disrupting the ability of P5 to change microtubule dynamics and growth cone morphology, as well as to reorganize cytoskeleton. *In vivo*, P5 accelerated cerebellum development in WT but not *clip1a* mutant zebrafish, and expression of exogenous CLIP1 in *clip1a* mutant promoted cerebellum development in response to P5. Thus, we have delineated the pathway by which P5 promotes cerebellum development by activating CLIP1 to promote microtubule dynamics leading to increased microtubule penetration into the growth cone and accelerated neurite outgrowth. This study reveals the mechanism by which P5 and CLIP1 function to promote neural development.

**Significance Statement:** *We have elucidated the mechanism of pregnenolone (P5) action:* P5 enhances brain functions, but its mode of action was unclear. Here we show that P5 activates CLIP1 to promote microtubule dynamics at the growth cone and to accelerate neural development.

*We have generated a zebrafish model of CLIP1 deficiency:* CLIP1 deficiency causes intellectual disability and defective neural development. Our zebrafish model can be used to study mechanisms related to this disease and other microtubule defects.

*We point to therapeutic intervention of neurological diseases using P5:* P5 is beneficial to the brain. We elucidate the mechanism of P5 action, thus accelerate the development of therapeutics using P5 and its derivatives.

## INTRODUCTION

### P5 is a neurosteroid present in nervous system

Pregnenolone (P5) is the first product of steroidogenesis catalyzed by CYP11A1 (cytochrome P450scc). It is present in the circulation in low amounts and is often considered as a mere precursor for all steroids, but it has a role in the brain. P5 is synthesized in neurons, astrocytes, oligodendrocytes and microglia (Mellon and Deschepper, 1993; Zwain and Yen, 1999). P5 level is altered in multiple sclerosis, Alzheimer’s disease and schizophrenia (Orefice et al., 2016), (Naylor et al., 2010; Cai et al., 2018).

P5 is a drug target for various brain diseases such as abuse-related disorders (Vallée, 2016; Tomaselli and Vallee, 2019), schizophrenia (Ritsner et al., 2014), and lower back pain (Naylor et al., 2020). P5 adjunct to risperidone attenuates core feature associated with autism spectrum disorders in clinical trials (Ayatollahi et al., 2020). In rodents, P5 improves learning and memory (Ducharme et al., 2010; Abdel-Hafiz et al., 2016), promotes recovery in a combination therapy after spinal cord injury of rats (Guth et al., 1994), and reduces synaptic defects associated with prenatal cannabis exposure (Frau et al., 2019). All these data point to the role of P5 in improving various brain functions.

### P5 targets regulate microtubule formation

P5 acts via a non-genomic mechanism (Weng and Chung, 2016). It has been associated with type 1 cannabinoid CB1 receptor (Vallee et al., 2014), a short-form of Shugoshin 1 (Hamasaki et al., 2014), estrogen receptor β (ERβ) (Shin et al., 2019), microtubule binding proteins CLIP1 (CLIP-170) (Weng et al., 2013) and MAP2 (Fontaine-Lenoir et al., 2006). Among these targets, CLIP1 stimulates microtubule polymerization (Weng et al., 2013), consistent with the role of P5 in promoting MT stability *in vivo* (Hsu et al., 2006). Microtubules are especially important for neurons. They participate in growth cone motility (Kahn and Baas, 2016). Loss of microtubules occurs in several neurodegenerative diseases (Zhang et al., 2015; Calogero et al., 2019). Impaired microtubule dynamics underlies neurodevelopmental disorders (Lasser et al., 2018; Barbiero et al., 2019).

CLIP1 is an microtubule plus-end binding protein (Yang et al., 2009; Nirschl et al., 2016). It forms a moving comet at the growing end of microtubules (Perez et al., 1999; Dixit et al., 2009), and regulates microtubule dynamics in growth cones (Neukirchen and Bradke, 2011). CLIP1 is involved in the pathogenesis of neurodevelopmental disorders (Barbiero et al., 2017), and *CLIP1* mutation causes intellectual disability in humans (Larti et al., 2015). P5 rescues neuronal defect in a CDKL5 deficiency caused by CLIP1 mislocalization (Barbiero et al., 2020).

CLIP1 activity is regulated by the conformational change. When its N- and C-termini interact with each other, CLIP1 assumes an inactive closed conformation (Lansbergen et al., 2004). Phosphorylation enhances this N- and C-terminal interaction leading to CLIP1 inactivation (Lee et al., 2010). The phosphomimic S311D mutant of CLIP1 is constitutively inactive, while S311A is constitutively active (Nakano et al., 2010). P5 interacts with CLIP1 at its central domain changing it into an active open conformation, and this activation is irrespective of the phosphorylation state of CLIP1 (Weng et al., 2013). But the role of CLIP1/P5 interaction in neurons is still unclear.

In this study we have analyzed P5 action in early stages of neuronal and cerebellum development. We show that P5 accelerates early neurite outgrowth, axon specification and elongation in several types of neurons. We show that P5 activates CLIP1 to promote cytoskeleton reorganization. *In vivo*, P5-CLIP1 also accelerates zebrafish cerebellum development. Thus, P5- CLIP1 promotes brain development by accelerating cytoskeleton maturation in developing neurons.

## Materials and Methods

### Plasmids and key resources

Plasmids, oligonucleotides, and other key resources are listed in Table 1. To generate pSC2+mOrange2-Flag-CLIP1, the mOrange2 region from pC1-mOrange2 was PCR amplified and ligated into the BamHI sites of the pSC2+-Flag-CLIP1 vector. The orientation of the insert was confirmed by BamHI/EcoRI digestion, followed by sequencing. The pDR274-pT7-Clip1a gRNA#2 plasmid was cloned by inserting annealed target oligonucleotides for gRNA#2 to pDR274 by *Bsa*I.

**Table 1.**
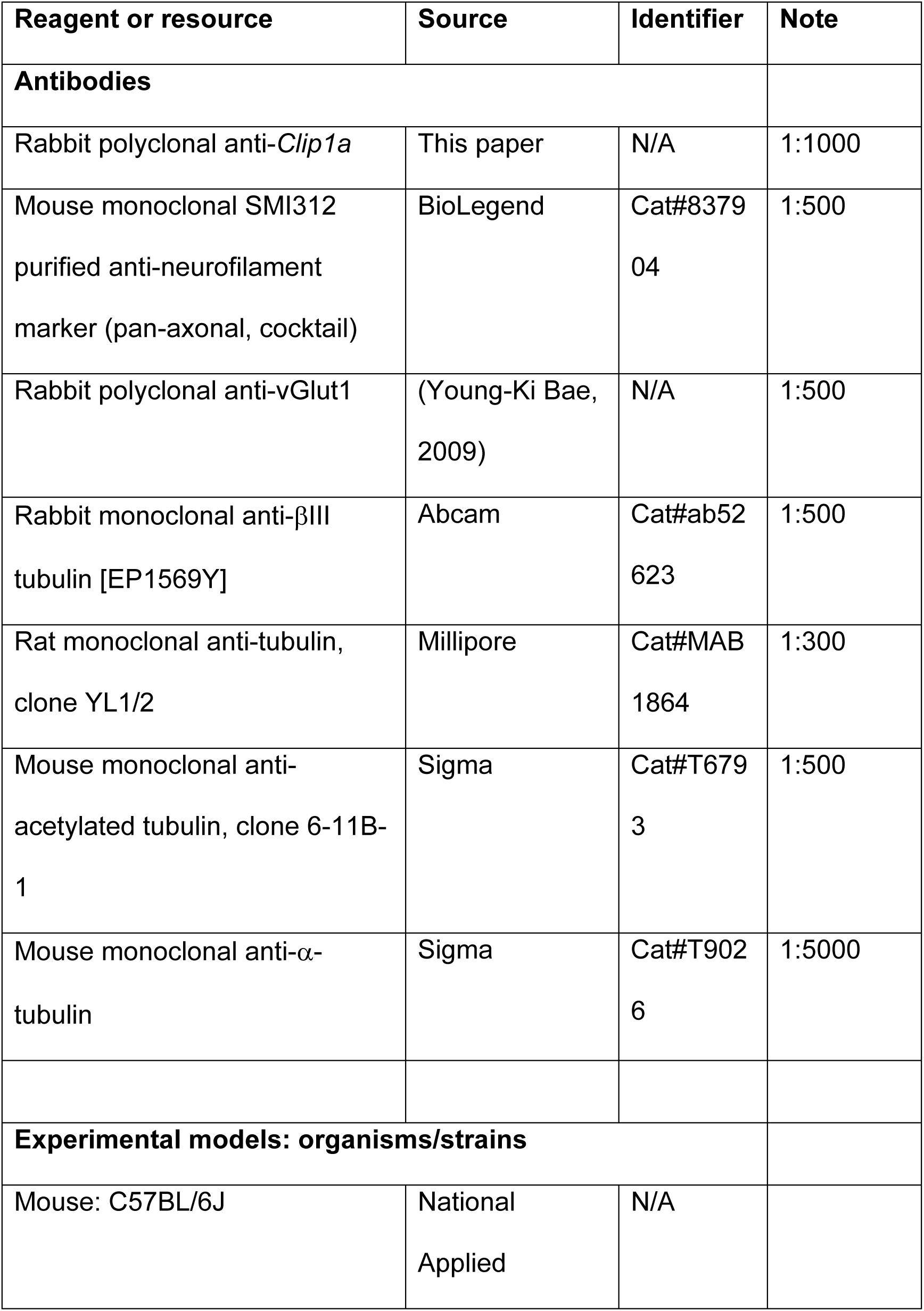
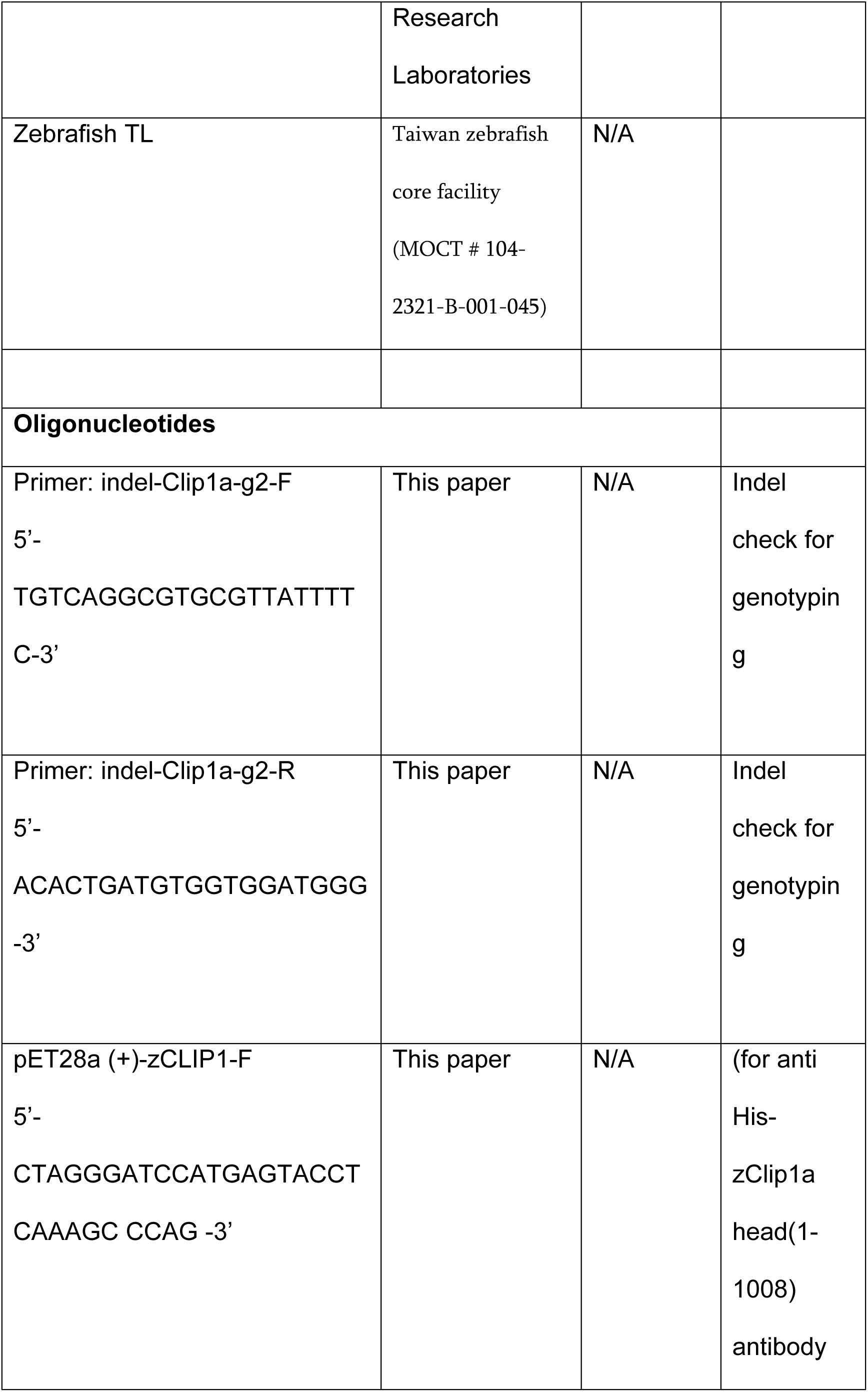
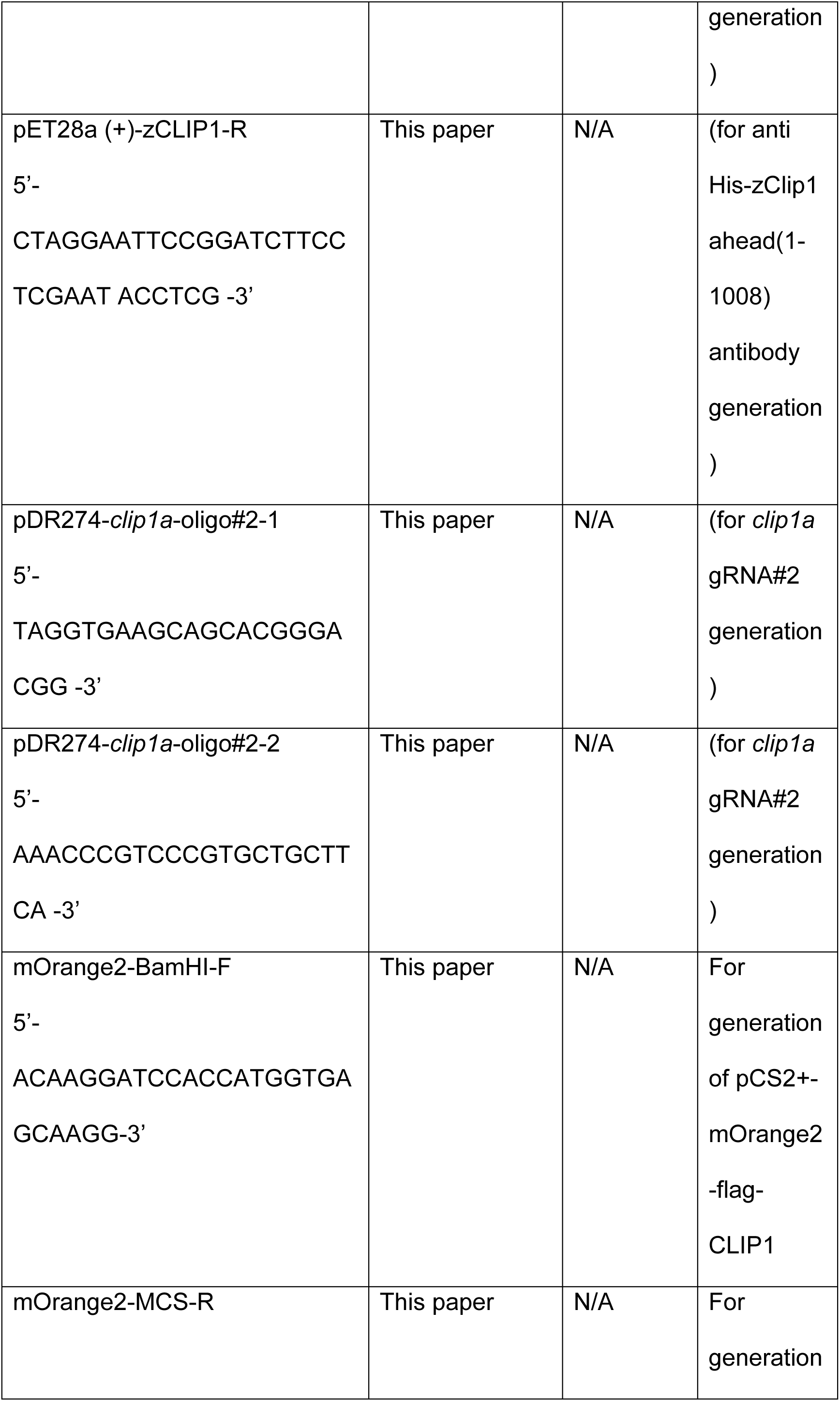
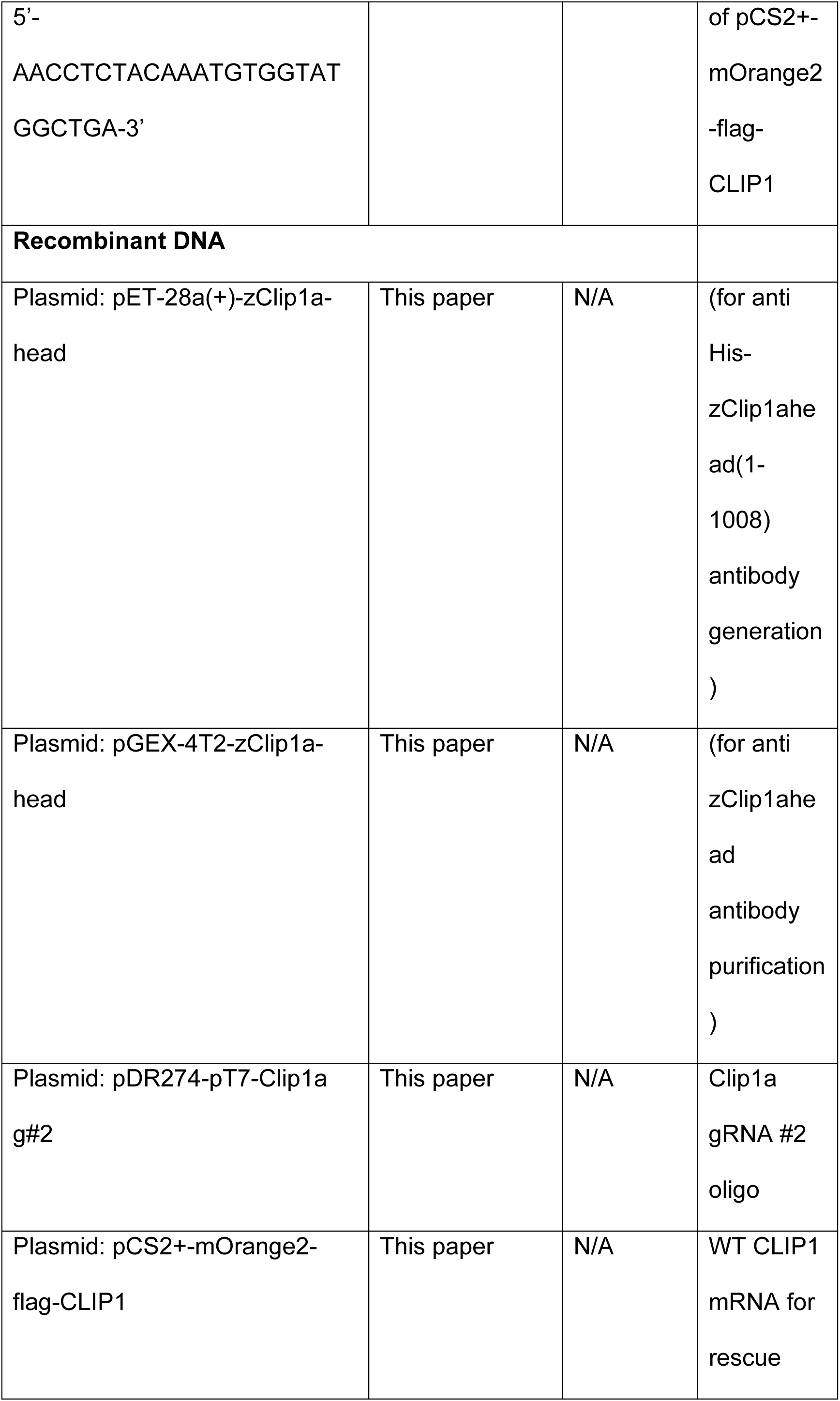
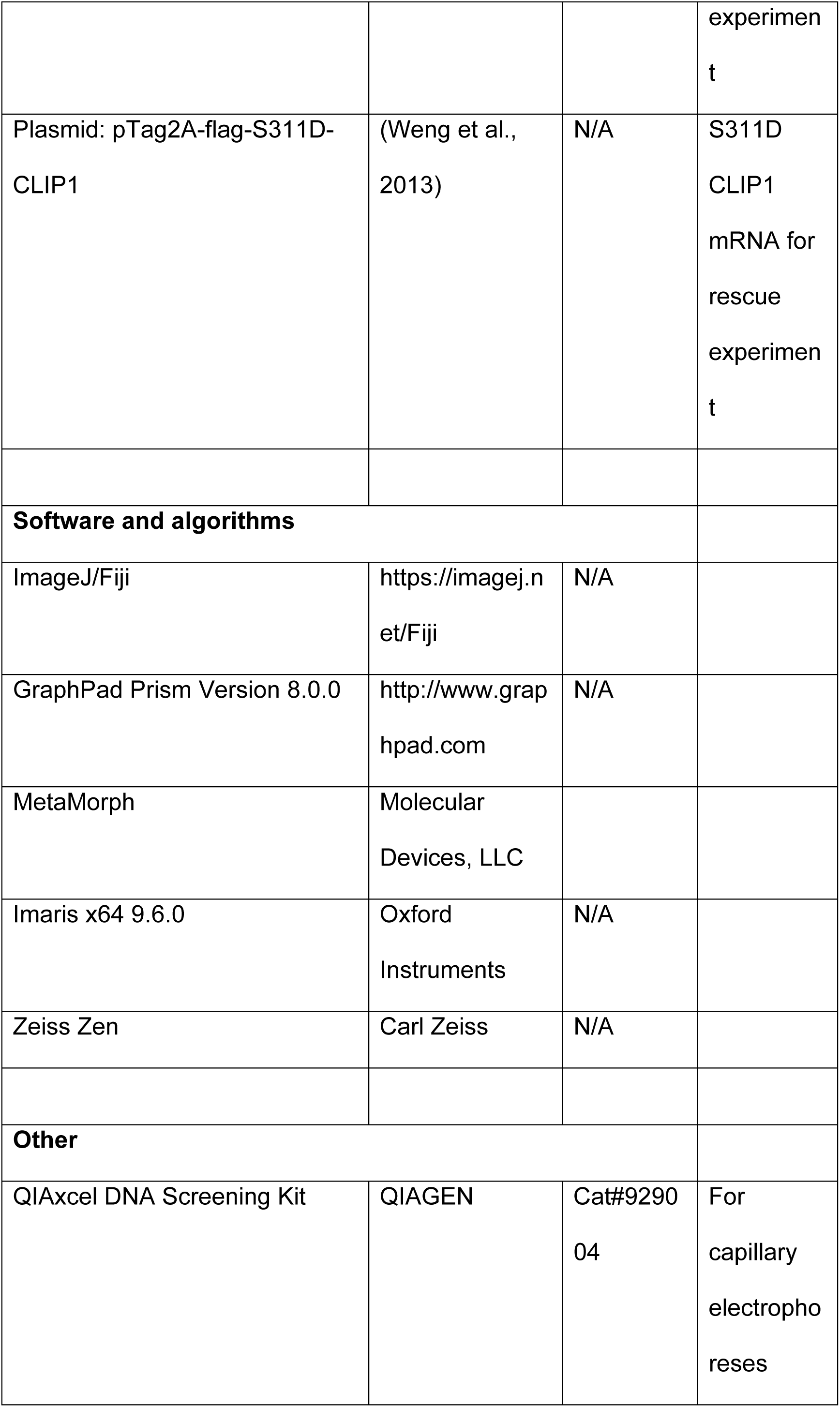
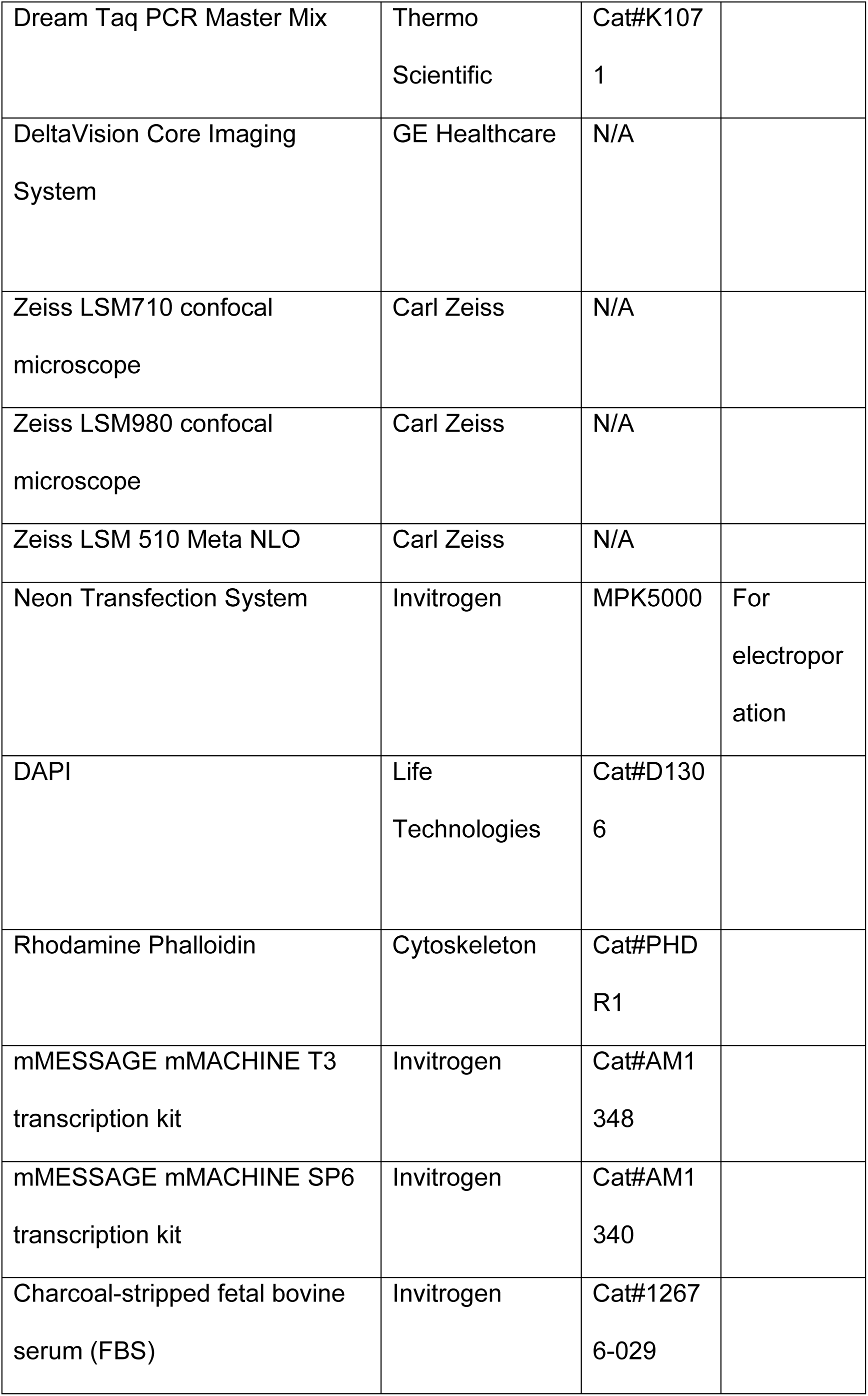
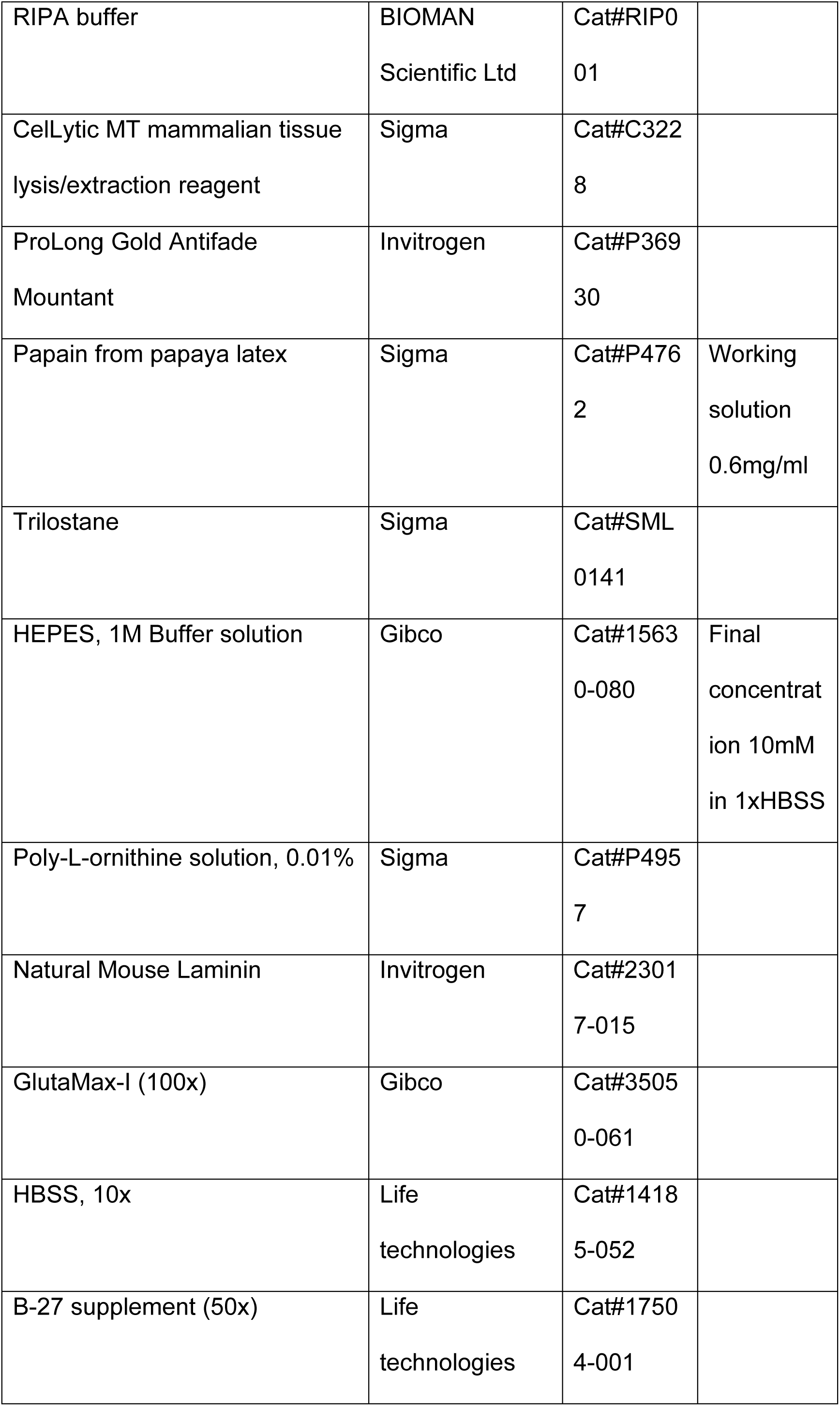
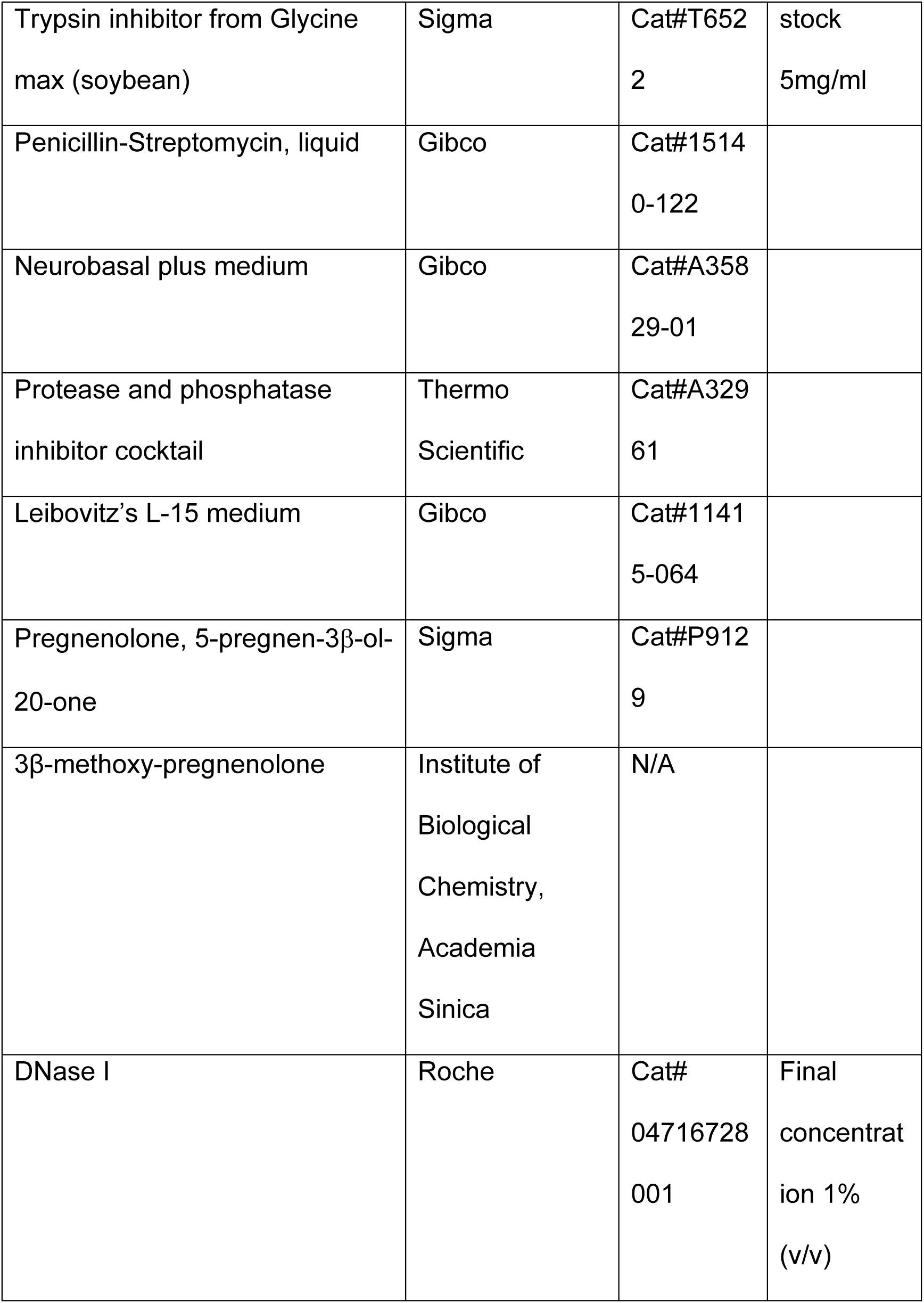
Key resources

### Animals

C57BL/6JNarl is a substrain of C57BL/6J provided by National Laboratory Animal Center of NARLabs, Taiwan (https://www.nlac.narl.org.tw/p3_animal2_detail.asp?cid=1&nid=3&ppage=0&type=1). All mice were maintained under temperature, light, humidity-controlled conditions, and 12h:12h light-dark cycle with free access to food and water and randomly assigned to experimental groups.

Zebrafish (Danio rerio, TL-strain) were reared at 28.5°C in 14:10 light-dark cycles according to the manual (Westerfield, 2000). Zebrafish sex cannot be determined until about two months post-fertilization, so their sex is unknown.

All animal work was approved by Institutional Animal Care and Utilization Committee of Academia Sinica, Taipei, Taiwan.

### Generation of zebrafish mutant line

Zebrafish *Clip1a* mutants were generated using the CRISPR-Cas9 technology. The gRNA sequences located at around AA 189 were designed and optimized using three different online tools. The sequence surrounding the gRNA site was examined in 24 fish to ascertain that there is no polymorphism in this specific region. For the generation of mutant fish, 100 pg gRNA and 200 pg Cas9 protein in 2 μl were co-injected into the yolk of zebrafish embryos before the 2-cell stage. The knockout efficiency was about 90% in the F0 founder. The founder fish were mated with wild type fish to generate F1 heterozygous mutant fish. The sequences of the F1 were examined and two heterozygous mutant fish with 4- and 8-bp deletion were kept for further breeding and experiments. Zebrafish genotypes were determined by incubating fin-clip lysate with indel-primers (Table 1) and DreamTaq PCR Master Mix (Thermo Scientific) at 95°C (3 min) of pre-denaturation and followed by 35 cycles of 95°C (30 s), 55°C (30 s), 72°C (30 s) and final extension 72°C (5 min). PCR products were analyzed by capillary electrophoresis with QIAxcel DNA Screening Kit according to the manufacturer’s instruction.

### Generation rabbit polyclonal antibody against zebrafish *Clip1a*

To raise rabbit polyclonal antibodies against zebrafish *Clip1a,* His-tag fusion (zClip1a-head) protein containing amino acids 1–336 of *Clip1a* was cloned into pET-28a(+) vector and the recombinant protein was overexpressed in *E. coli* BL21DE3. The His-tag fusion protein was purified and used for immunization. The resulting rabbit serum purified by binding to antigen-bound PVDF membrane and eluted with 500 μl buffer (5 mM glycine, 150 mM NaCl pH2.4). The purified antibody was neutralized with 100 μl 0.1 M phosphate buffer pH8.0.

### Primary neuronal culture

Mouse hippocampal and cortical neuronal cultures were prepared from mice from both sexes at postnatal day 0 as described previously (Beaudoin et al., 2012). After dissection in cold Hank’s buffered salt solution (1xHBSS,10mM HEPES), meninges were removed and mouse cortex or hippocampus were digested with Papain working solution (0.6 mg/ml Papain, 0.5 mM EDTA, 0.5 mM CaCl2, 0.2 mg/ml L-cysteine in 1x HBSS) containing DNase I for 20 min at 37°C, washed in 1xHBSS three times and manually triturated in Neurobasal plus medium. Neurons were counted, resuspended with plating media (Neurobasal plus medium, B-27 supplement, 2 mM GlutaMax-I, 0.45 % sucrose, Penicillin-Streptomycin) and 10^5^ cell/cm^2^ plated on plastic dish or glass coverslips coated with Poly-L-ornithine/laminin.

For cerebellar granule neuron culture, cerebellum was dissected from mice at postnatal day 5-6, in HBSS, digested with 0.05%Trypsin-EDTA, containing DNase I 15 min at 37°C. soybean trypsin inhibitor (0.5 mg/ml) and 0.4 % BSA in HBSS was added to dissociation mixture and incubated 1 min 37°C, then cooled on ice and triturated with a fire-polished glass Pasteur pipette. Cell suspension washed in HHBS with 0.5% BSA, resuspended in HHBS, overlaid on Percoll solution (40% Percoll, 1xPBS, 0.2% glucose, 2 mM EDTA), and centrifugated at 2600 rpm at 4°C for 10 min, and resulting pellet of cerebellar granule neurons was washed with HHBS,0.5%BSA. Cells were resuspended with plating media (Neurobasal plus media, B-27 supplement, 2 mM GlutaMax-I, 250 µM KCl, 0.45% sucrose, Penicillin-Streptomycin) and 10^5^ cell/cm^2^ plated on plastic dish or glass coverslips coated with Poly-L-ornithine/laminin.

Zebrafish embryonic forebrain neuronal culture was prepared as described previously (Chen Z. 2013). Zebrafish embryos from 42 hpf were dechorionated, decapitated in E3 embryo media (5 mM NaCl, 0.17 mM KCl, 0.33 mM CaCl2, 0.33 mM MgSO4, 0.00001% Methylene Blue) with 0.04% (m/v) tricaine methanesulfonate (MS-222). Forebrain was isolated by custom- made fine tungsten filaments and incubated in 0.05% EDTA-Trypsin 15 min at RT, centrifuge shortly to remove supernatant and triturate in equal volume of Leibovitz’s L-15 medium containing 2% (v/v) charcoal-stripped fetal bovine serum (FBS), 0.4% (v/v) Penicillin-Streptomycin. Cell suspension was plated on plastic dishes or glass coverslips coated with Poly-L-ornithine solution, followed by laminin. Culture was maintained at RT during 7 h in Leibovitz’s L- 15 medium containing B-27 supplement (100x), 1 mM GlutaMax-I, 0.1% (v/v) Penicillin-Streptomycin. The fine tungsten filaments for dissection were prepared according to instructions from former members of Dr. John R. Henley’s lab, and its tips were attached to 9V battery lead (positive and negative) and opposite tips were placed in saturated sodium hydroxide solution to complete the circuit during 2-3 h.

### Whole embryo immunochemistry

For immunohistochemistry embryos were treated with 0.005% phenylthiourea to prevent the development of pigmentation as described previously (Young-Ki Bae, 2009). The larvae were fixed at 4°C in 4% PFA in PBST (PBS, 0.1% Triton X-100) for 3 h. The fixed larvae were washed with PBST, and incubated in acetone at − 20 °C for 15 min. Larvae were washed once with PBST and twice with PBS-DT (PBS, 1% BSA, 1% DMSO, 1% Triton X-100), and incubated in 5% normal goat serum, PBS-DT at room temperature for 1 h. The samples were incubated with vGLUT1 antibody solution (1/500) at 4°C overnight. After four washes with PBST, the samples were incubated with secondary antibodies (1/1000 dilution, Alexa Fluor 555 goat anti-rabbit IgG (H + L), Invitrogen) for 2h at room temperature, following three washes with PBST. Nuclei were counterstained with DAPI.

The whole-mount immunostaining of zebrafish larvae with anti- acetylated tubulin antibody was performed as described previously (Jian Zhu, 2020). The larvae were fixed with 2% trichloroacetic acid in PBS for 3 h at room temperature, followed by three washes with PBS-T (PBS, 0.8% Triton X- 100) for 10 min each. Then larvae were dehydrated and rehydrated through graded methanol concentration (50% MeOH once, 100% twice, and 50% once, 10 min each), and rinsed three times in PBS-T for 5 min each. The larvae were treated by ice-cold acetone for 20 min at –20 °C, rinsed three times in PBS-T. Fish were digested with 10 ng/µl proteinase K for permeabilization, re-fixed with fresh 4% PFA in PBS for 20 min at RT, rinsed three times in PBS-T for 10 min each, and blocked with 10% normal goat serum PBS-TD (PBS, 1% DMSO, 0.8% Triton X-100) for 3 h at RT, followed by incubation with primary antibody (1:500) overnight at 4°C. After three rinses in PBS-T for 30 min each, embryos were incubated with secondary antibodies (1:1000 dilution, Alexa Fluor 555 goat anti-mouse IgG (H + L), Invitrogen) for 3h at RT, followed by three PBS-T washes, 5 min DAPI staining and two more PBS-T washes.

Fluorescence images of stained embryos were acquired using an LSM- 710 confocal microscope, and Z-series stacks were used for 3D analysis by Imaris x64.

### Immunoblotting. (WB for *zClip1a* antibody)

Fish embryos at 24 hpf (hours postfertilization) were homogenized with a hand pestle (Toyobo) in cell lysis buffer containing protease and phosphatase inhibitor cocktail on ice. The lysates were centrifuged at 13,000 rpm for 20 min at 4°C. Protein (10 µg protein per lane) was separated by SDS polyacrylamide gel, transferred to a PVDF membrane (Immobilon, Millipore). The membranes were blocked for 1 h in TBS-TW buffer (50 mM Tris/HCl, pH 7.5, 150 mM NaCl, and 0.1% Tween-20) containing 5% BSA and then incubated overnight at 4°C with purified rabbit polyclonal anti-Clip1a (1:1000) or mouse monoclonal anti-α-tubulin antibodies (1:5000). Immunoblots were developed using horseradish peroxidase-conjugated goat anti-mouse or goat anti-rabbit IgG antibodies, followed by chemiluminescence system (Millipore).

### Immunocytochemistry

For immunocytochemistry all primary neuronal cultures were plated on glass coverslips coated with poly-L-ornithine and mouse laminin.

The mouse primary neuronal culture were rinsed with pre-warmed at 37°C PBS and fixed with fresh-prepared 4%PFA/4% sucrose in PBS for 10 min. Fixed cells were permeabilized with 0.1% Triton X-100 in PBS for 5 min, blocked with 5% normal goat serum/1% BSA in PBS-Tw (0.05% Tween-20) 1 h at RT and incubated in 2.5% normal goat serum/1% BSA in PBS-Tw overnight at 4°C with the primary antibodies listed in Key resource table.

The primary culture of zebrafish forebrain neuron was fixed with 4%PFA/0.01% glutaraldehyde in cytoskeleton stabilizing buffer (60 mM PIPES, 25 mM HEPES, 5 mM EGTA, 1 mM MgCl, pH 7.0) (Witte H.2008) for 20 min, permeabilized with 0.1% Triton X-100 in PBS 3 min and blocked with 5% normal goat serum/1% BSA in PBS-Tw 30 minutes at room temperature. Samples were incubated in 5% normal goat serum/1% BSA in PBS-Tw overnight at 4°C with primary antibody listed in Table 1.

All samples were incubated with Alexa-dye labeled secondary antibodies (1/4000) in 2.5% normal goat serum/1% BSA in PBS-Tw 1.5 h at room temperature. F-actin was stained in fixed cells with Phalloidin-TRITC (Sigma-Aldrich) according to the manufacturer’s instruction. Nuclei were stained with DAPI. The samples were mounted using Prolong Gold Antifade (Molecular Probes).

### *In vitro* transcription

RNAs were synthesized by reverse-transcription from pCS2+mOrange2-Flag-CLIP1 or pTag2A-flag-S311D-CLIP1 plasmids using mMESSAGE mMACHINE *in vivo* transcription kits according to manufacturer’s instructions. For rescue experiments, 150 pg synthesized RNAs were injected into zebrafish embryos at the one-cell stage.

### Transfection

The purified cerebellar granule neurons were transfected with pSC2+mOrange2-Flag-CLIP1 by electroporation with Neon Transfection System. according manufacturer instruction and plated on a glass cover slips coated with poly-L-ornithine, laminin. The live-cell imaging for microtubule comet tracking was performed at DIV 2.

### Drug treatment

Pregnenolone in 10 mM DMSO stock solutions were diluted by Neurobasal or Leibovitz’s L-15 media to 1 mM working solution and used for preparation of medium aliquot with final drug concentration depending on total volume of plating media in each well. Volume of drug containing medium aliquot did not exceed 10% of total volume of medium in each well and was added after removing equal volume of plating medium from well to avoid osmotic shock. Final DMSO concentration was 0.01% or less. Drugs were added to the cell cultures 1 h after plating and maintained through the experiment.

Zebrafish eggs were collected 10 minutes after spawning and maintained in a Petri dish with filtered drug-containing medium. Zebrafish medium was exchanged every day.

### Microscope data acquisition and image analysis

For all neurites/axon outgrowth analysis phase contrast images were collected with LCM 510 Meta NLO microscope using an LD Plan-Neofluar 40×/0.6 Ph2 Corr objective. Cells in randomly chosen areas in each well were counted. During acquisition live primary neuronal culture plated on plastic dishes were maintained within 37°C temperature, humidity, and 5 % CO2 control specimen incubator.

Fluorescently labeled neurons were imaged with LSM710, individual growth cone for filopodia analysis were imaged with LSM980 Airyscan (Zeiss), equipped with a Plan-Apochromat 63×/1.40 Oil DIC M27 objective.

MetaMorph (Molecular Devices, LLC) was used for neurite/axon outgrowth, tubulin PTM and F-actin/tubulin colocalization analysis in developing neurons. All neurites and axon were manually traced, following extraction of data files for statistical analysis. The threshold criteria for definition of live polarized neurons were described previously (Yamamoto et al., 2012). The difference in lengths of longest neurite from average length of sibling minor processes >10 μm is required to induce the polarized state of neurons. This criterium was used to exclude non-polarized neurons from axon outgrowth analysis. To analyze the ratio of tubulin PTM fluorescent intensity of ROI from IIIβ-tubulin channel was used to generate mask, applied to acetylated tubulin and tyrosinated tubulin individual channels images. The area of F-actin and IIIβ-tubulin colocalization in growth cone was analyzed by Measure colocalization instrument of MetaMorph. Fluorescent intensity profiles for filopodia were generated by the image analysis module Zen 9.0 (Zeiss). All data were extracted for statistical analysis by GraphPad.

Zebrafish larval cerebellum was imaged using an LSM710 laser scanning confocal microscopy with a C-Apochromat 40x/1.2 W Corr objective (Zeiss) generating a z-stack (step size 1µm). Imaris X64 9.6.0 (Oxford Instruments) was used to analyze total volume of fluorescently labeled 3D cerebellum. Threshold setting was kept the same throughout the whole experiment. The shape and borders of developing cerebellum were defined by DAPI staining.

### Microtubule comet measurement

The live-cell imaging of mOrange2-Flag-CLIP1 comets in cerebellar granule neurons was performed in a temperature and humidity-controlled chamber using a DeltaVision Core fluorescence microscope (GE Healthcare) with a 60x U-APO objective (Nikon). As transfected CLIP-170 is prone to aggregate into strongly fluorescing immobile puncta, all immobile puncta were excluded from the quantification. Images were taken at 2 second intervals for 1 minute and deconvoluted with Applied Precision softWorx imaging software (GE Healthcare). The microtubule comet tracking and quantification was performed with the MTrackJ plugin in ImageJ/Fiji (Meijering et al., 2012). To quantify comet characteristics, individual comet locations were manually marked for each imaging time point. MTrackJ calculates the lifetime, speed, and extension length of each comet, among other characteristics.

### Experimental Design and Statistical Analysis

All experiments were independently repeated for at least 3 or more times, and data are expressed as means ± standard deviation (SD) to show the variability. One-Way ANOVA was used for all statistical analysis when comparing more than two groups of samples. Two-tailed Student’s *t* test was used when comparing two groups of samples. All statistics were performed using GraphPad Prism software.

## Results

### Pregnenolone increases early neurite outgrowth in different types of neurons

To examine the effect of P5, we cultured primary neurons that extend neurites in bimodal or multi-modal neuron polarity (Fig. 1A). In bipolar systems such as cerebellar granule neurons and zebrafish forebrain neurons, cells initially form a single neurite and later develop a second neurite at the opposite side. In multipolar models such as hippocampal and cortical neurons, cells grow several neurites before the longest neurite becomes the primary axon (Tahirovic and Bradke, 2009).

**Fig. 1.**
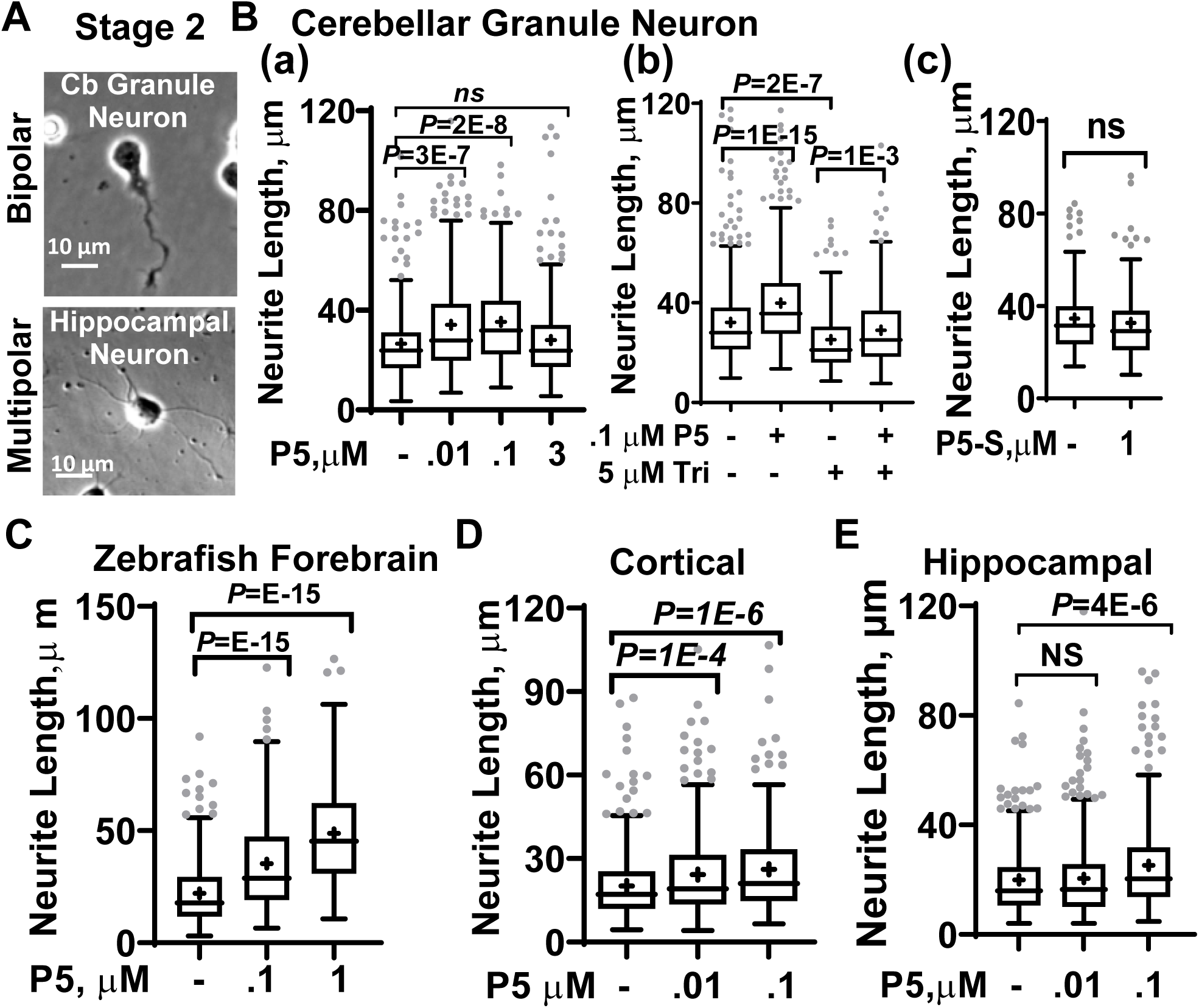
P5 accelerates early neurites outgrowth. **A.** Images of cerebellar (Cb) granule neurons (top) and hippocampal neurons (bottom) at stage 2 of development (24 h after plating). **B. (a)** P5 promotes neurite lengths in cerebellar granule neurons at 0.01 and 0.1 µM, F(3,1267)=17.36, *p*=4*E- 11, one-way ANOVA. (**b)** even in the presence of steroidogenesis inhibitor, trilostane (Tri). F(3,1664)=55.0, *p*=1*E-15, one-way ANOVA. (**c)** P5 sulfate (P5-S) did not affect neurite length of cerebellar granule neurons. t(291)=1.05, *p*=0.29, unpaired *t.* test. P5 at 0.1 µM and 1 µM increased neurites outgrowth in. **C**. zebrafish forebrain, F(2,832)=30.5, *p*=1*E-13, one-way ANOVA; **D**. cortical F(2,994)=13.4, *p*=1*E-6, one-way ANOVA and **E**. hippocampal neurons F(2,1243)=14.1, *p*=8*E-7, one-way ANOVA. Data are represented as Tukey’s box-and-whiskers plots with the mean values shown as the “+” sign.

We examined primary neuronal cultures at stage 2 of development, when differentiating neurons start to extend neurites. The neurites of mouse cerebellar granule neurons were 27 ± 15 µm; they grew to 34 ± 20 µm when treated with 0.01 µM P5 and 35 ± 18 µm at 0.1 µM P5 (*p*=4*E-11, one-way ANOVA, Fig. 1Bab). However, neurites remained the same length (28 ± 17 µm) when cells were treated with 3 µM P5, suggesting the effective concentration range for P5 is narrow.

To ascertain that we examined the effect of P5 but not its metabolites, we treated cells with trilostane, which inhibits the conversion of P5 to progesterone (P4) and all subsequent steroids (Young et al., 1996). P5 (0.1 µM) still promoted neurite outgrowth from 25 ± 13 µm to 29 ± 15 µm in the presence of 5 µM trilostane (*p*=1*E-15, one-way ANOVA, Fig. 1Bb). Another P5 metabolite, pregnenolone sulfate, did not affect neurite length (34 ± 14 µm vs. 33 ± 15 µm, *p*=0.2, *t* test, Fig. 1Bc). Thus P5, but not its metabolites, promotes neurite outgrowth of differentiating cerebellar granule neurons.

We also examined the effect of P5 in other differentiating neurons, including the zebrafish forebrain neuron that extend a single neurite and neurons that extend multiple neurites such as the cortical neuron and hippocampal neuron. In zebrafish forebrain neurons, neurite length increased from 22 ± 14 µm to 35 ± 21 µm at 0.1 µM P5 and 49 ± 23 µm at 1 µM P5 (*p*=1*E-13, one-way ANOVA, Fig. 1C). In cortical neurons, neurite lengths were increased from 20 ± 12 µm to 26 ± 16 µm when cells were treated with 0.01 µM P5 and to 24 ± 16 µm with 0.1 µM P5 (*p*=1*E-6, one-way ANOVA, Fig. 1D). In hippocampal neurons, neurites grew from 20 ± 14 µm to 25 ± 18 µm at 0.1 µM P5, but remained 20 ± 14 µm at 0.01 µM P5 (*p*=8*E-7, one-way ANOVA, Fig. 1E). Our data thus show that P5 accelerates neurite outgrowth in four different types of differentiating neurons in a dose-dependent manner. The range of effective P5 concentration is the same among different type of neurons and independent from their polarity types.

### Pregnenolone accelerates axon specification and increases axon outgrowth

Hippocampal neurons at Stage 3 (DIV 2-3) develop nascent axons from the longest neurite (Tahirovic and Bradke, 2009). To analyze axon outgrowth, we stained Stage-3 (DIV-2) hippocampal neurons with IIIβ-tubulin and F-actin (Fig. 2A). The crosslinked microtubule bundles appeared as a line, while fast-growing disorganized microtubules appeared like gaps along the axonal shaft (Fig. 2Aa). The balance between the gap and the microtubule bundle is important for axon homeostasis (Hahn et al., 2019). The length of the gap was increased from 44 ± 12% of the total length to 59 ± 9% when neurons were treated with 1 µM P5 (*p*=7*E-3, *t* test Fig. 2Ab), but the number of gaps was not changed (3.2 ± 1.6 vs. 3.7 ± 2.6, *p*=0.6, *t* test, Fig. 2Ac). These data show that P5 shifts the balance of microtubule bundles and gaps into the fast-growing microtubules along the axonal shaft.

**Fig. 2.**
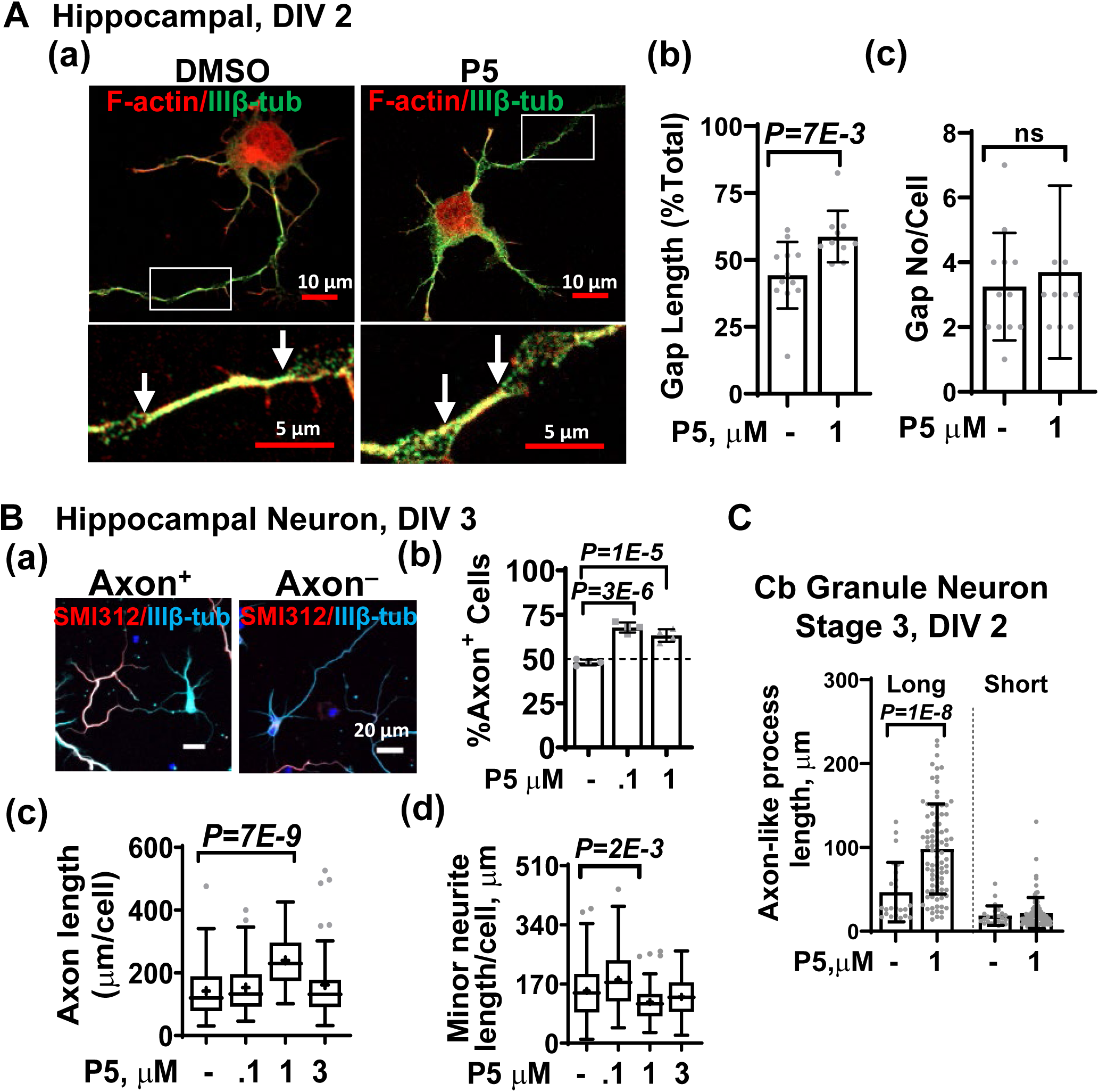
P5 promotes axon development. **A.** P5 accelerates hippocampal axon growth. (**a**) DIV2 hippocampal neurons immunostained with F-actin (red) and IIIβ-tubulin (green). The boxed regions are enlarged below, showing microtubule bundles as lines demarcated by white arrows and bulged “gaps”. **(b)** P5 increases the percentage of dynamic “gap” fragments length t(20)=3.0, *p*=0.007, unpaired *t test*; **(c)** but not the numbers of “gaps” t(20)=0.48, *p*=0.63, unpaired *t test*. **B.** P5 promotes hippocampal axon development at DIV 3. (**a)** Hippocampal neurons immunostained with anti-SMI312 (red) and anti-IIIβ-tubulin (blue) antibodies. **(b)** P5 increases the percentage of hippocampal neurons with specified axon F(2,9)=55.5, *p*=8*E-6, one-way ANOVA. (**c)** Axon length per cell is increased by 1 µM P5, F(3,333)=13.1, *p*=3*E-8, one-way ANOVA, and **(d)** minor neurite length increased by 0.1 µM P5, F(3,333)=9.8, *p*=3*E-6, one-way ANOVA. **C.** P5 promotes the growth of long axon-like processes at stage 3 of cerebellar (Cb) granule neuronal culture (DIV2), but has no effect on short neurites F(3,212)=65.9, *p*=1*E- 15, one-way ANOVA.

We examined axon specification of hippocampal neurons using a pan-axonal marker SMI-312 (Fig. 2Ba) (Wilkins et al., 2003). About 48 ± 2 % hippocampal neurons already specified their axons, and this number increased to 68 ± 3 % and 63 ± 4 % when cells were treated with 0.1 µM and 1 µM P5, respectively (*p*=8*E-6, one-way ANOVA, Fig. 2Bb). The lengths of the nascent axons were increased from ± 91 to 240 ± 83 µm when treated with 1 µM P5, but remained as 153 ± 80 µm and 162 ± 115 µm when cells were treated with 0.1 µM and 3 µM P5, respectively (*p*=3*E-8, one-way ANOVA, Fig. 2Bc). Lower concentration of P5 (0.1 µM) increased the lengths of minor neurites from 150 ± 81 µm to 182 ± 80 µm, but not higher concentrations of 1 µM and 3 µM P5 (118 ± 57 µm and 132 ± 58 µm, respectively, *p*=3*E-6, one-way ANOVA, Fig. 2Bd). This result indicates that P5 functions in a narrow concentration range; it loses functions when its concentration is too high.

Mouse cerebellar granule neurons develop two axon-like processes at Stage 3 (DIV 2). The long process will become an axon, while the short neurite will retract and be replaced by dendrites in later stages (Tahirovic and Bradke, 2009). We examined the development of these axon-like processes. The length of the long axon-like process was increased from 47 ± 35 µm to 98 ± 53 µm when cells were treated with1 µM P5, but short process was not affected (19 ± 11 µm vs. 22 ± 18 µm, *p*=1*E-15, one-way ANOVA, Fig. 2C). Thus, P5 accelerates axon formation for bipolar cerebellar granule neurons.

### Pregnenolone increases dynamic microtubules and accelerates microtubule stability along nascent axon

Neurons have compartment-specific distribution of tubulin post-translational modifications that affects microtubule dynamics (Song and Brady, 2015). To differentiate types of tubulin post-translational modifications we performed immunocytochemistry of growing cerebellar granule neurons with antibodies specific to tyrosinated (Tyr), acetylated (Ac) and IIIβ-tubulin, representing dynamic, stable, and total microtubules, respectively (Fig. 3A). In growing axonal-like processes of cerebellar granule neurons, The dynamic Tyr-tub^+^ pools of microtubules were increased from 0.31 ± 0.06 to 0.39 ± 0.11 when cells were treated with 1 µM P5 (*p*=8*E-5, *t* test), but Ac-tub^+^ stable microtubules pools were not affected (0.5 ± 0.13 vs 0.5 ± 0.16, *p*=0.9, *t* test) (Fig. 3B).

**Fig. 3.**
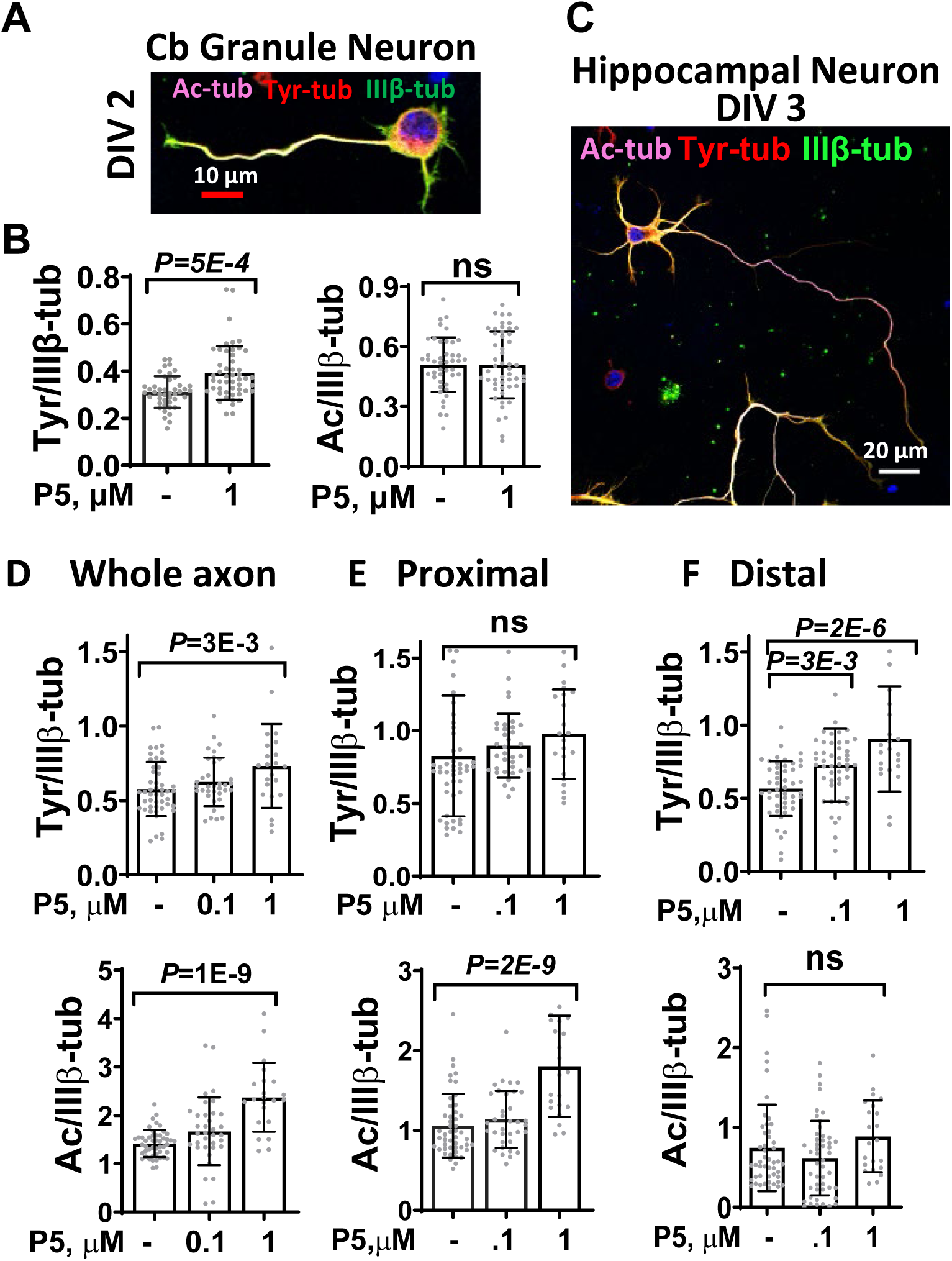
P5 regulates post-translational modifications of tubulin in developing neurons. **A-B.** P5 increases dynamic MT pool in developing axon-like processes of cerebellum granule neurons at DIV2. **A**. Immunostaining of cerebellum granule neurons with anti-acetylated (pink), anti-tyrosinated (red) and anti-IIIβ-tubulin (green) antibodies. **B**. P5 promotes tyrosination but not acetylation of tubulin along the axon-like processes. **C**. Hippocampal neurons immunostained with anti-acetylated (pink), anti-tyrosinated (red) and anti-IIIβ-tubulin (green) antibodies. P5 increases tubulin acetylation and tyrosination in the (**D)** whole axon, but increases only acetylation in the **(E)** proximal axon, and increases tyrosination only in the (**F)** distal axon.

We also examined tubulin modifications in the developing axon of DIV3 hippocampal neurons (Fig. 3C). P5 (1 µM) increased dynamic microtubules from 0.6 ± 0.2 to 0.7± 0.3, (*p*=3*E-3, one-way ANOVA) and stable microtubule pools from 1.4 ± 0.3 to 2.4 ± 0.7, (*p*=1*E-9, one-way ANOVA) in the whole axon (Fig. 3D). On closer examination, in the proximal 15-μm segment, where axon starts to form, P5 did not affect dynamic Tyr-tub (0.8 ± 0.4 vs 1.0 ± 0.3, *p*=0.09, one-way ANOVA), but increased the level of acetylated tubulin from 1.1 ± 0.4 to 1.8 ± 0.7 (*p*=2*E-9, one-way ANOVA) (Fig. 3E). At the distal site close to the growth cone, dynamic Tyr-tub^+^ microtubules were increased from 0.56 ± 0.18 to 0.73 ± 0.24 when treated with 0.1 µM P5 (*p*=2*E-3), and to 0.9 ± 0.4 with 1 µM P5 (*p*=7*E-7, one-way ANOVA), but Ac-tub^+^ stable microtubules were not changed by 1 µM P5 (0.7 ± 0.5 vs 0.9 ± 0.4, *p*=0.26, one-way ANOVA) (Fig. 3F). These data indicate that P5 effect is compartment-specific and P5 initially increases dynamic microtubule pool.

P5 accelerate microtubule dynamics in non-neuronal cell by activating its receptor CLIP1 (Weng et al., 2013). To measure microtubule dynamics, we transfected cerebellar granule neurons with CLIP1 fused with the fluorescent protein mOrange (Fig. 4A). We tracked fluorescent CLIP1-mOrange comets located at the tip of the growing microtubules and measured their lifetime, polymerization distance, and speed. P5 (1 µM) prolonged comet lifetime from 6.4 ± 1.7 s to 7.4 ± 1.8 s (*p*=3*E-2, *t* test, Fig. 4B) and increased the comet track length from 2.4 ± 0.8 µm to 3.3 ± 1.3 µm (*p*=1*E-3, *t* test, Fig. 4C), resulting in faster comet polymerization speed from 0.39 ± 0.11 µm/s to 0.45 ± 0.14 µm/s) (*p*=3*E-2, *t* test, Fig. 4D). These results indicate that P5 promotes CLIP1 attachment to the microtubule and increased microtubule dynamic in developing neurons.

**Fig. 4.**
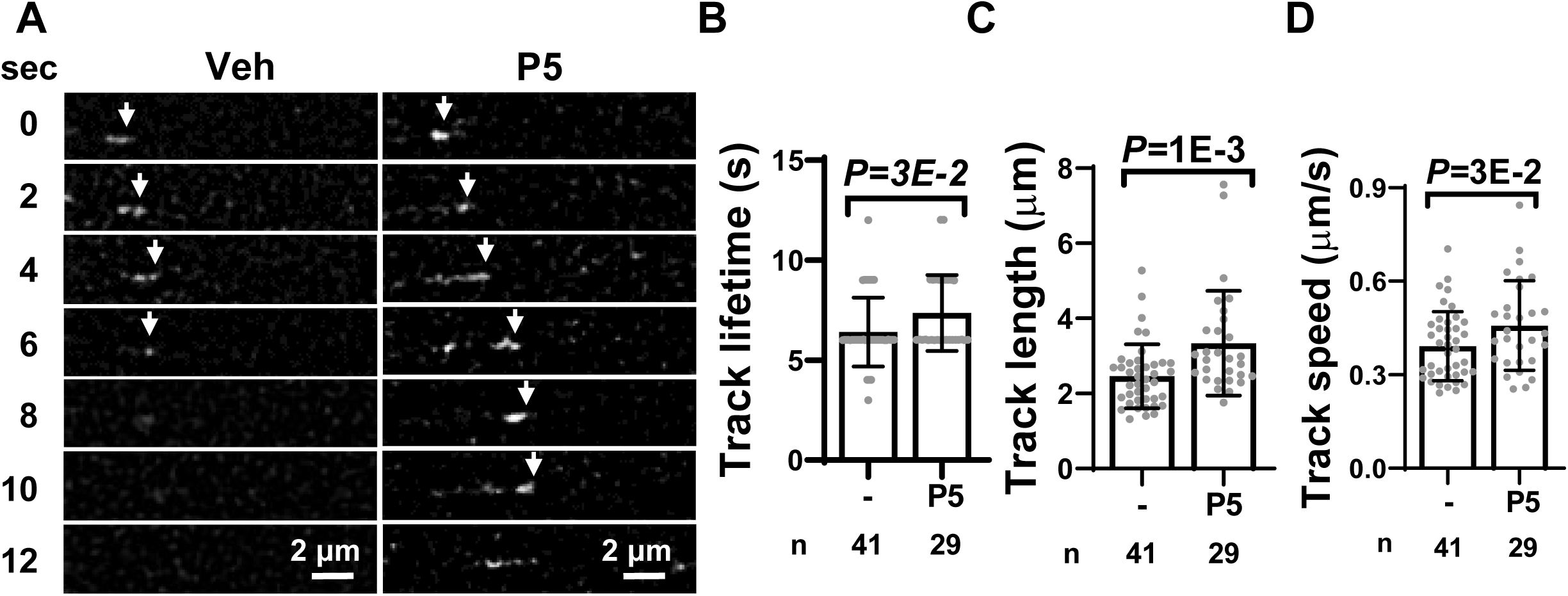
P5 accelerates movement of microtubule comets in cerebellar granule neurons. **A.** time lapse imaging of microtubule comets. White arrowheads point to the comet in question. P5 (1 µM) increases **(B)** comet track life times, t(68)=2.19, *p*=0.03, unpaired *t* test, (**C**) comet track lengths, t(68)=3.27, *p*=0.001, unpaired *t* test and (**D**) comet track speed t(68)=2.18, *p*=0.03, unpaired *t* test.

### CLIP1 mediates pregnenolone effect to promote neurite outgrowth

To investigate the role of P5 receptor CLIP1 in developing neurons, we generated *clip1a* knockout zebrafish lines by CRISPR/Cas9. Two knockout lines were used in the study. These two lines harbor 4-bp (*clip1a^4d^*) and 8-bp (*clip1a^8d^*) deletions (Fig. 5A), which could be differentiated by capillary electrophoresis (Fig. 5B). These mutations lead to premature protein termination at 189 and 154 AA, respectively (Fig. 5C). To detect zebrafish Clip1, we generated antibodies against zebrafish Clip1a. This antibody detected full-length 170-kDa Clip1a in the 24-hpf wildtype embryos (Fig. 5D). Both *clip1a^4d^* and *clip1a^8d^* fish, however, did not produce Clip1a detected by immunoblotting (Fig. 5D). Therefore, mutations in these two lines are completely null.

**Fig. 5.**
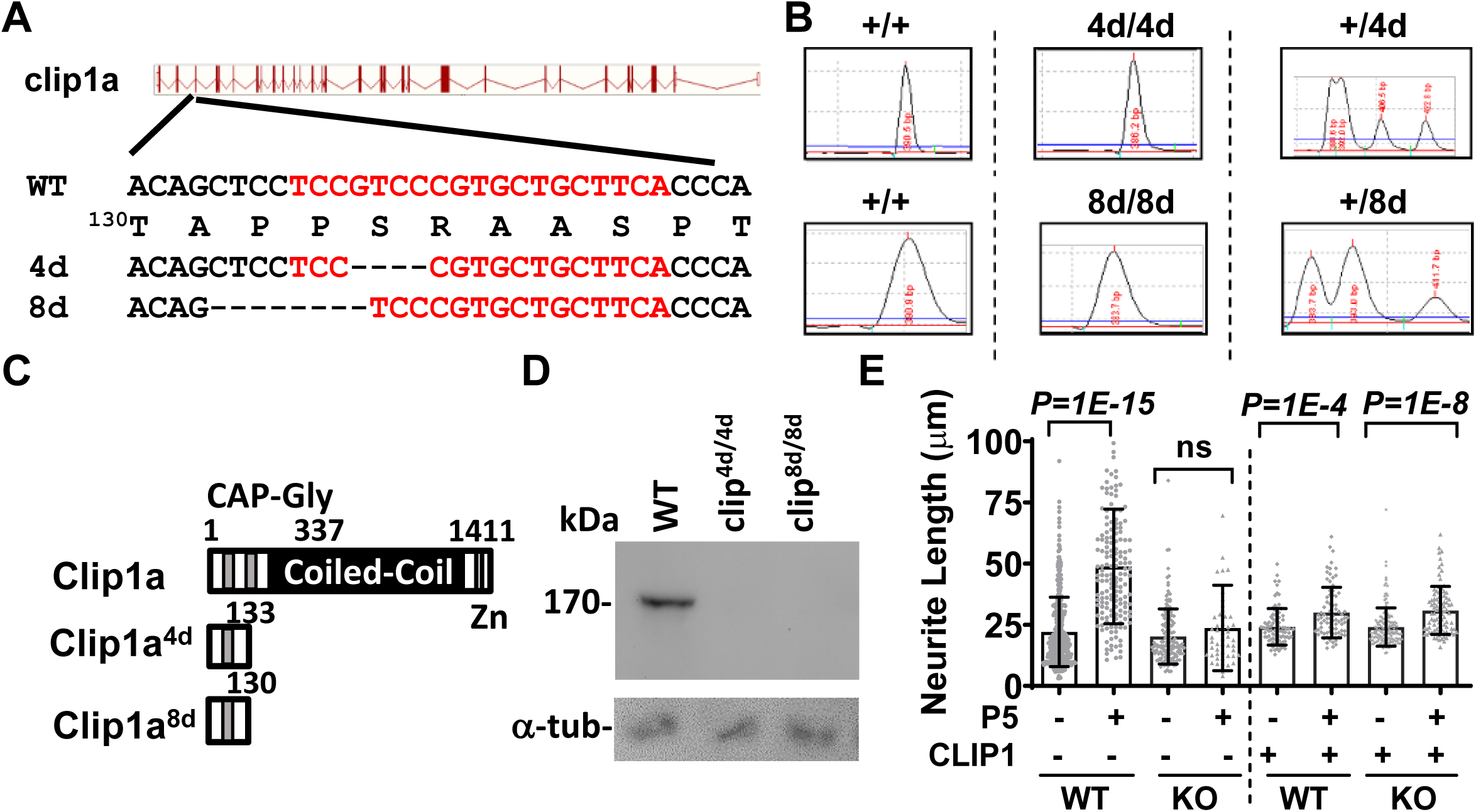
P5 promotes outgrowth of neurites from zebrafish forebrain culture by activating CLIP1. **A** The genomic structure of zebrafish *clip1a* showing exons as boxes and introns as gaps. The coding sequences of WT and mutant (4d and 8d) alleles in exon 3 are shown with the gRNA sequences marked in red. **B.** Genotyping of WT and mutant alleles by capillary gel electrophoresis. The assigned genotypes are written on top of the DNA tracings. **C.** The domain structures of zebrafish wildtype and mutant Clip1a. The CAP-Gly domains are marked by grey boxes, and the coiled-coil domain marked by a black box. Two mutants, Clip1a^4d^ and Clip1a^8d^ are predicted to be truncated at 189 and 154 AA, respectively. **D**. Immunoblots detects 170-kDa zebrafish Clip1a in 24-hpf zebrafish embryos but not in the *Clip^4d/4d^* and *clip^8d/8d^*mutants. **E**. P5 promotes outgrowth of neurites from zebrafish forebrain neurons but not of *Clip1a* KO (*clip^8d/8d^*) neurites. Dashed line separates data from un-injected (left panel) and wild type *CLIP1* mRNA injected (right panel) embryos. *p*=1*E-15, one-way ANOVA. Data are presented as mean±SD.

Primary neuronal culture was prepared from 42-hpf forebrain of zebrafish embryos, and the outgrowth of their neurites was analyzed. P5 (1 µM) increased the length of WT neurites derived from 22 ± 14 µm to 49 ± 23 µm, but not of *clip1a^8d^/^8d^* knockout (KO) neurites (20 ± 11 µm vs. 24 ± 17 µm) (Fig. 5E, left panel). To validate the role of CLIP1, we forced CLIP1 production in neurons by injecting human *CLIP1* mRNA into one-cell stage zebrafish embryo before preparing primary neuronal culture. Upon expressing CLIP1, the neurite lengths increased from 24 ± 7 µm to 30 ± 10 µm by P5 in WT neurons, and from 24 ± 7 µm to 31 ± 9 µm in *clip1a^8d^/^8d^* neurons (Fig. 5E, right panel). The rescue of P5 effect by CLIP1 demonstrates that P5 increases neurite outgrowth via CLIP1.

### CLIP1 mediates pregnenolone effect to balance dynamic and stable microtubule pools

We then examined whether the effects of P5 on microtubules depend on CLIP1. Cultured zebrafish forebrain neurons were stained with antibodies against Tyr-, Ac-, and IIIβ-tubulin to differentiate dynamic and stable microtubules (Fig. 6A). P5 (1 µM) increased dynamic tyrosinated tubulin along the WT neurite from 0.9 ± 0.6 to 1.5 ± 1.1 (*p*=1*E-3, one-way ANOVA), but not the *clip1a^8d/8d^*homozygous (HO) knockout neurites (0.8 ± 0.4 vs 0.9 ± 0.5, *p*=0.6, one-way ANOVA) (top panel, Fig. 6B). The injection of *CLIP1* mRNA rendered *clip1a* KO neurons responsive to P5, and tyrosinated tubulin pool was now increased by 1 µM P5 from 0.9 ± 0.5 to 1.2 ± 0.5, *p*=1*E-3, one-way ANOVA) (top panel, Fig. 6C). But CLIP1 overexpression abolished P5 effect on tyrosinated tubulin in WT neurons (0.7 ± 0.4 vs 0.9 ± 0.4, *p*=0.06, one-way ANOVA), while enabling P5 to increase the amount of stable Ac-tubulin from 1.2 ± 0.7 to 1.8 ± 1.3, *p*=1*E-2, one-way ANOVA) (bottom panel, Fig. 6C). This result indicates that too much activated CLIP1 increases stable but not dynamic microtubules.

**Fig. 6.**
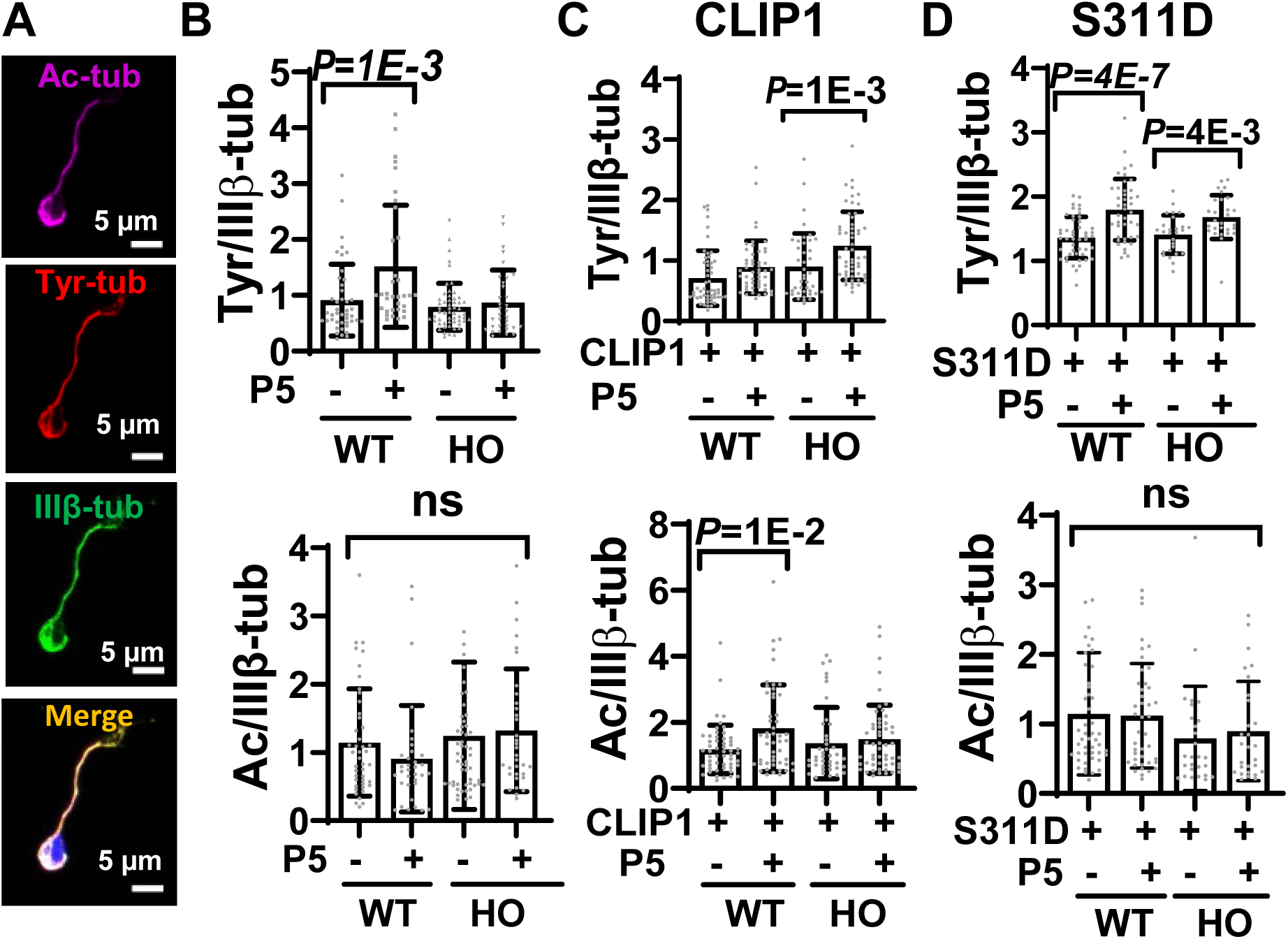
P5 shifts equilibrium of dynamic/stable microtubule pools towards microtubule network maturation via CLIP1. **A**. Immunostaining of zebrafish primary forebrain neurons with anti-acetylated (pink), anti-tyrosinated (red), IIIβ-tubulin (green) antibodies. (**B**) Quantitation shows that 1 µM P5 increases tyrosinated tubulin in the neurites of primary forebrain neurons prepared from WT embryos, but not from *clip1a^8/8d^* homozygous (HO) mutant embryos F(3,162)=8.6, *p*=2*E-5, one-way ANOVA. Acetylated tubulin is not affected by P5 in both WT and HO primary forebrain neurons, F(3,162)=1.36, *p*=0.25, one-way ANOVA. (**C**) *CLIP1* mRNA injection confers increased tyrosinated tubulin of HO mutant in response to P5, but depletes the tyrosinated tubulin response of WT neurites F(3,194)=10.6, *p*=1*E-6, one-way ANOVA. CLIP1, instead, increases acetylated tubulin pool of WT neurites in response to P5 F(3,194)=3.28, *p*=2*E-2, one-way ANOVA. (**D**) S311D-CLIP1 increases tyrosinated tubulin of both WT and HO neurites F(3,140)=12.43, *p*=2*E-7, one-way ANOVA, but does not change acetylated tubulin F(3,140)=1.57, *p*=0.19.

The S311D-CLIP1 has closed conformation that can be activated by P5 (Lee et al., 2010). The overexpression of S311D-CLIP1 potentiated P5 effects to increase dynamic Tyr-tubulin pools from 1.4 ± 0.3 to 1.8 ± 0.4 (*p*=4*E-7, one-way ANOVA) in WT neurons, and from 1.4 ± 0.2 to 1.7 ± 0.3 (*p*=8*E-3, one-way ANOVA) in and HO mutant culture (top panel, Fig. 6D). But it had no effect on the stable Ac-tubulin pools in both WT (1.1 ± 0.8 vs 1.1 ± 0.7, *p*=0.8, one-way ANOVA) and HO mutant (0.8 ± 0.7 vs 0.9 ± 0.7, *p*=0.6, one-way ANOVA) (bottom panel, Fig. 6D).

Summarizing all these data, we show that P5 regulates the balance of tubulin post-translational modifications in developing neurons by activating CLIP1 in a dose-sensitive manner.

### Pregnenolone regulates cytoskeleton reorganization in growth cone of developing neurons via CLIP1

CLIP1 regulates microtubule dynamic in growth cone (Neukirchen and Bradke, 2011). To analyze the effect of P5 on cytoskeleton organization in growth cone we stained F-actin with Rhodamine-Phalloidin and total tubulin with IIIβ-tubulin antibody in mouse cerebellar granule neurons at 48h after plating. The growth cones of cerebellar granule neurons initially spread out, but became narrower upon P5 treatment (Fig. 7Aa). When quantified, the total area decreased from 119 ± 84 µm^2^ to 69 ± 50 µm^2^ when treated with 0.1 µM P5, (*p*=8*E-5), and to 73 ± 59 µm^2^ with 1 µM P5 (*p*=1*E-5, one-way ANOVA) (Fig. 7Ab). The number of filopodia in the growth cone was decreased from 0.9 ± 0.2 to 0.6 ± 0.2 when cells were treated with 0.1 µM P5 (*p*=5*E-7), and to 0.7 ± 0.3 with 1 µM P5 (*p*=1*E-4, one-way ANOVA) (Fig. 7Ac). The transition zone is the region where microtubules overlaps with F-actin in the growth cone. This zone was increased from 34 ± 20% to 65 ± 24% when cells were treated with 0.1 µM P5, (*p*=1*E-13) and to 52 ± 22% with 1 µM P5 (*p*=1*E-7, one-way ANOVA) (Fig. 7Ad).

**Fig. 7.**
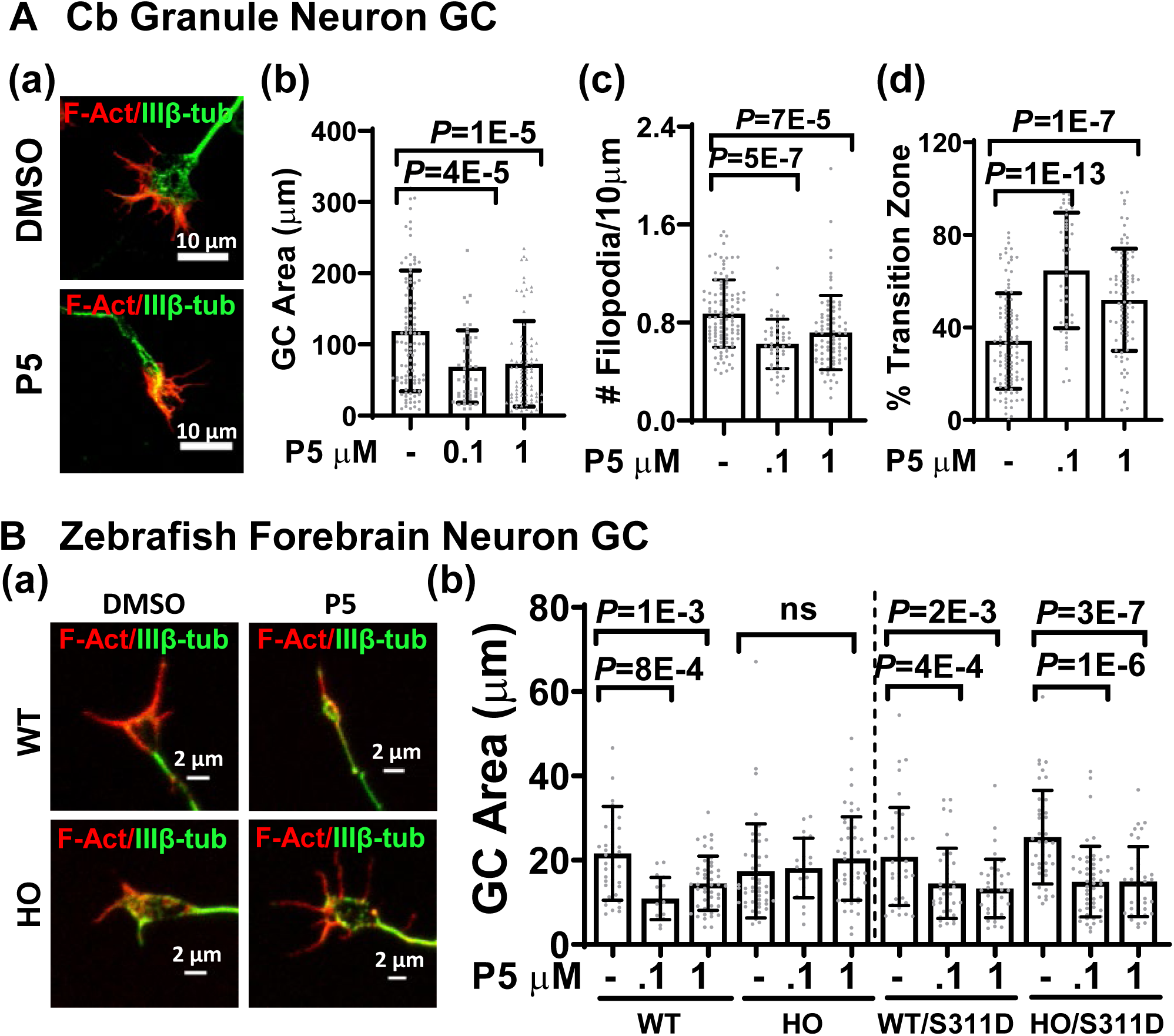
P5 regulates growth cone (GC) morphology. **A.** P5 changes GC morphology of cerebellar granule neurons. **a**. Immunostaining of GC by F-actin (red) and IIIβ-tubulin (green). Quantitation of GC area covered by F-actin and microtubules show that P5 decreases (**b)** GC area, *p*=4*E-6, and **(c)** decreases numbers of filopodia, *p*=4*E-7, but **(d)** increases transition zone (F-Act^+^IIIβ-tub^+^ region) in GC of cerebellar granule neurons, *p*=7*E- 4,. **B**. P5 changes GC morphology of zebrafish forebrain neurons via CLIP1. **(a)** Immunostaining of F-actin (red), IIIβ-tubulin (green) of zebrafish forebrain neurons; **(b)** P5 decreases total area of growth cone in WT but not in *Clip1a* KO neurons. Expression of S311D-CLIP1 renders HO mutants to decrease GC area in response to P5, *p*=1*E-11. Data are presented as mean±SD.

To test whether CLIP1 is involved in this P5 effect, we stained zebrafish forebrain neurons with Rhodamine-Phalloidin for F-actin and with IIIβ-tubulin antibody for total microtubules (Fig. 7Ba). The area of the WT growth cone was decreased from 22 ± 11 µm^2^ to 11 ± 4 µm^2^ when cells were treated with 0.1 µM P5 (*p*=6*E-4), and to 14 ± 6 µm^2^ with 1 µM P5 (*p*=3*E-4, one-way ANOVA). It was not changed in *clip1a* homozygous knockout HO neurons (17 ± 11 µm^2^ vs 18 ± 7 µm^2^ at 0.1 µM P5, *p*=0.7, and vs 20 ± 9 µm^3^ at 1 µM P5, *p*=0.1, one-way ANOVA) (Fig. 7Bb). Injection of *S311D-CLIP1* mRNA encoding phosphomimic CLIP1 enabled P5 to decrease area of WT growth cone from 21 ± 11 µm^2^ to 14 ± 8 µm^2^ when cells were treated with 0.1 µM P5 (*p*=2*E-3), and to 13 ± 6 µm^2^ with 1 µM P5 (*p*=3*E-4), one-way ANOVA) (Fig. 7Bb). Similarly, upon S311D overexpression, The area of HO growth cones decreased from 25 ± 11 µm^2^ to 15 ± 8 µm^2^ when cells were treated with at 0.1 µM P5 (*p*=8*E-8) and to 15 ± 8 µm^2^ with 1 µM P5 (*p*=7*E-7, one-way ANOVA) (Fig. 7Bb). These results demonstrate the P5 activates CLIP1 to change growth cone morphology and reduces its areas.

To understand the role of P5 in the growth cone, we examined cytoskeleton structures there in more detail. The filopodia at the edge of the growth cone are rich in F-Actin, but contain very little microtubules. Upon P5 treatment, microtubules penetrated deeply into the filopodium tips and this microtubule contain tyrosinated tubulin (Fig. 8Aa, white arrowheads). We traced and quantified the fluorescence intensities of F-actin, Tyr-tub and IIIβ- tub along the filopodia (Fig. 8Ab). Tyr-tub in the distal half of the filopodia was increased from 0.18 ± 0.09 to 0.7 ± 0.6 when cells were treated with 1 µM P5 (p=1*E-15, *t* test, Fig. 8Ac) and total IIIβ-tubulin increased from 0.18 ± 0.10 to 0.57 ± 0.3 (*p*=1*E-15, *t* test, Fig. 8Ad). However, the distribution of F-actin at the distal and proximal part of filopodia was the same with or without P5 treatment (0.6 ± 0.3 vs 0.6 ± 0.4, *p*=1*E-15, *t* test, Fig. 8Ae).

**Fig. 8.**
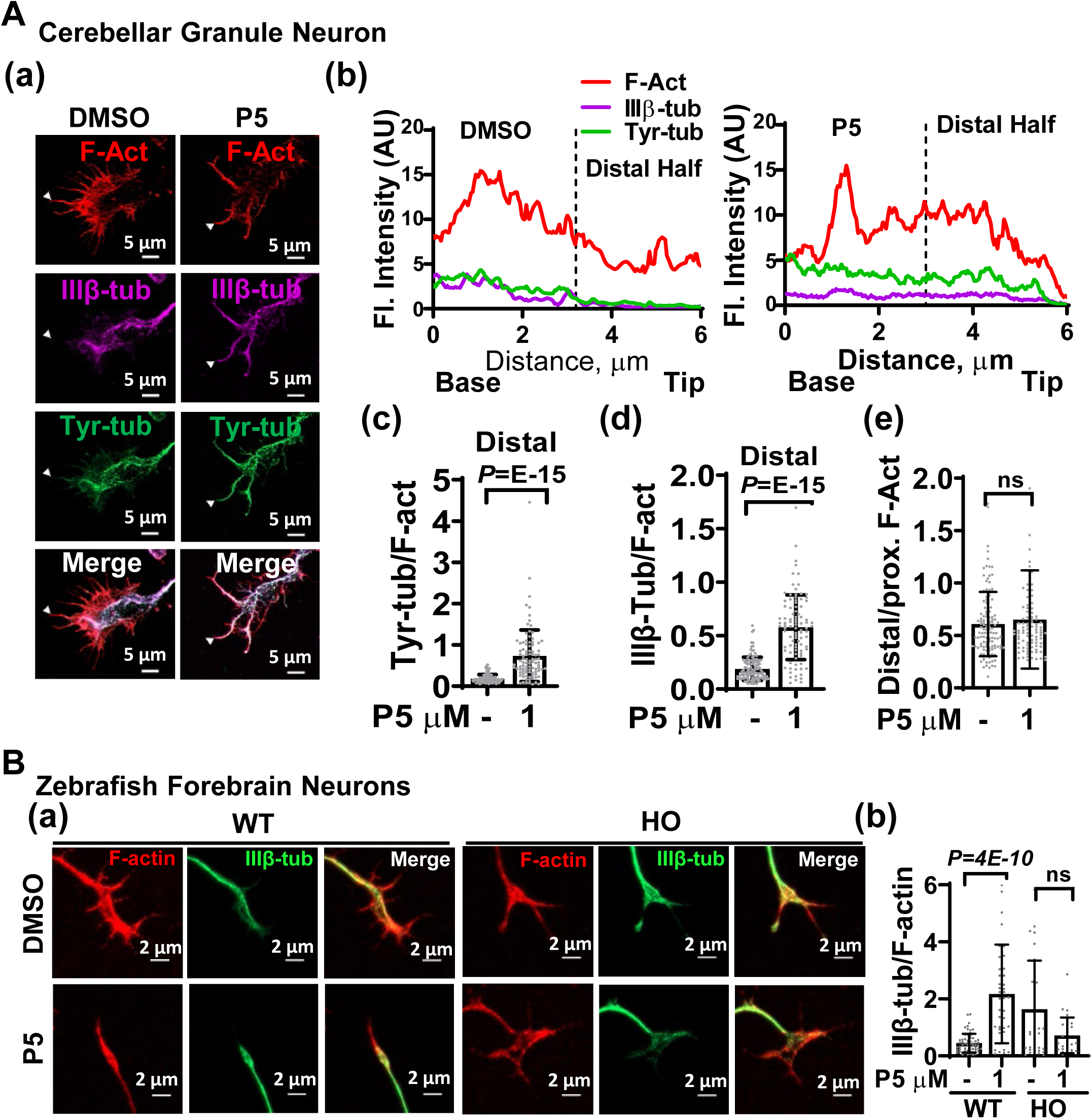
P5 modulates cytoskeleton reorganization in filopodia. **A.** P5 promotes dynamic MT invasion into the distal half of filopodia in cerebellar granule neurons; **a**. GC of cerebellar granule neurons immunostaining of F-actin (red), IIIβ-tubulin (pink), tyrosinated tubulin (green). White arrowheads point to individual filopodia, traced in (b). **b.** Traces of cytoskeleton profiles from the base to the tip of filopodia. P5 increases **(c)** tyr-tubulin t(214)=9.56, *p*=1*E-15, unpaired *t* test, and **(d)** total IIIβ-tubulin t(214)=13.2, *p*=1*E-15, unpaired *t* test vs. F-actin in the distal half of filopodia. **(e)** P5 does not affect the distribution of F-actin along the filopodia t(214)=0.82, *p*=0.4, unpaired *t* test. **B. (a)** GC of WT and homozygous (HO) mutant zebrafish forebrain neurons immunostained with F-actin (red), IIIβ-tubulin (green). **(b)** P5 increases the penetration of tubulin into F-actin area in the distal half of filopodia of wild type but not *clip1a* HO neurons F(3,176)=41.74, *p*=4*E-9, one-way ANOVA. Data are presented as mean±SD.

We also examined cytoskeleton distribution in the growth cone of zebrafish forebrain neuronal culture (Fig. 8Ba). In the WT growth cone, microtubule penetration into the filopodia was increased from 0.4 ± 0.3 to 2.2 ± 1.7 when cells were treated with 1 µM P5 (*p*=2*E-9, one-way ANOVA), but not in *clip1a* knockout filopodia (1.6 ± 1.7 vs 0.7 ± 0.6 at 1 µM P5, *p*=0.6, one-way ANOVA, Fig. 8Bb). These data demonstrate that P5 increases microtubule penetration into the filopodial tip of the growth cone via activating CLIP1.

### Pregnenolone accelerates zebrafish cerebellum development via CLIP1

In addition to the primary neuronal culture, we investigated the role of P5 *in vivo*. We analyzed axons in zebrafish embryos at 2.5 dpf and 3 dpf, when cerebellum starts to form (Kani et al., 2010). Total volume of stable microtubule tracks in the developing cerebellar axons was detected by immunostaining with anti-acetylated tubulin antibody (Fig. 9A) and quantified. P5 increased the volume of developing microtubule tracks in the WT neurons increased from 1.9 ± 0.7 x10^-4^ µm^3^ to 2.7 ± 0.6 x10^-4^ µm^3^ when treated with 1 µM P5 (*p*=2*E-4, one-way ANOVA), and from and (2.4 ± 0.5 x10^-4^ µm^3^ to 3.0 ± 0.8 x10^-4^ µm^3^ (*p*=1*E-2, one-way ANOVA) in the *clip1a* heterozygous (HET) neurons at 2.5 dpf (Fig. 9Ba). But the volume in the *clip1a* homozygous HO mutants was not affected by P5 (2.2 ± 0.7 x10^-4^ µm^3^ vs 2.4 ± 0.5 x10^-4^ µm^3^, *p*=0.3, one-way ANOVA, Fig. 9B). Similarly, at 3 dpf the volume of developing microtubule tracks increased from 4.6 ± 1.3 x10^-4^ µm^3^ to 7 ± 3 x10^-4^ µm^3^ (*p*=1*E-2, one-way ANOVA) in the WT neurons and from 5.3 ± 2.8 x10^-4^ µm^3^ to 7.5 ± 2.2 x10^-4^ µm^3^ (*p*=2*E-2, one-way ANOVA) in *clip1a* heterozygous (HET) embryos. But it was not changed in the *clip1a* homozygous HO mutants (4.8 ± 3.0 x10^-4^ µm^3^ vs 5 ± 3 x10^-4^ µm^3^, *p*=0.8, one-way ANOVA, Fig. 9B). We also analyzed functional axons in the zebrafish cerebellum by staining with anti-vGlut1 antibody, a marker of presynaptic terminal of cerebellar granule neurons (Bae et al., 2009). the volume of functional axon in 3-dpf cerebellum was increased by P5 from 1.6 ± 0.8 x10^-4^ µm^3^ to 2.9 ± 1.3 x10^-4^ µm^3^ (*p*=1*E-4, one-way ANOVA) when WT embryos were treated with 1 µM P5, and from 2.3 ± 0.9 x10^-4^ µm^3^ to 3.3 ± 0.9 x10^-4^ µm^3^ (*p*=2*E-2, one-way ANOVA) in *clip1a* HET (Fig. 9D). But P5 did not change the axon volume of *clip1a* HO knockout cerebellum (3.3 ± 1.2 x10^-4^ µm^3^ vs 2.7 ± 1.4 x10^-4^ µm^3^, *p*=0.12, one-way ANOVA, Fig. 9D). These data demonstrate that P5 accelerates axon formation during cerebellum development via CLIP1.

**Fig. 9.**
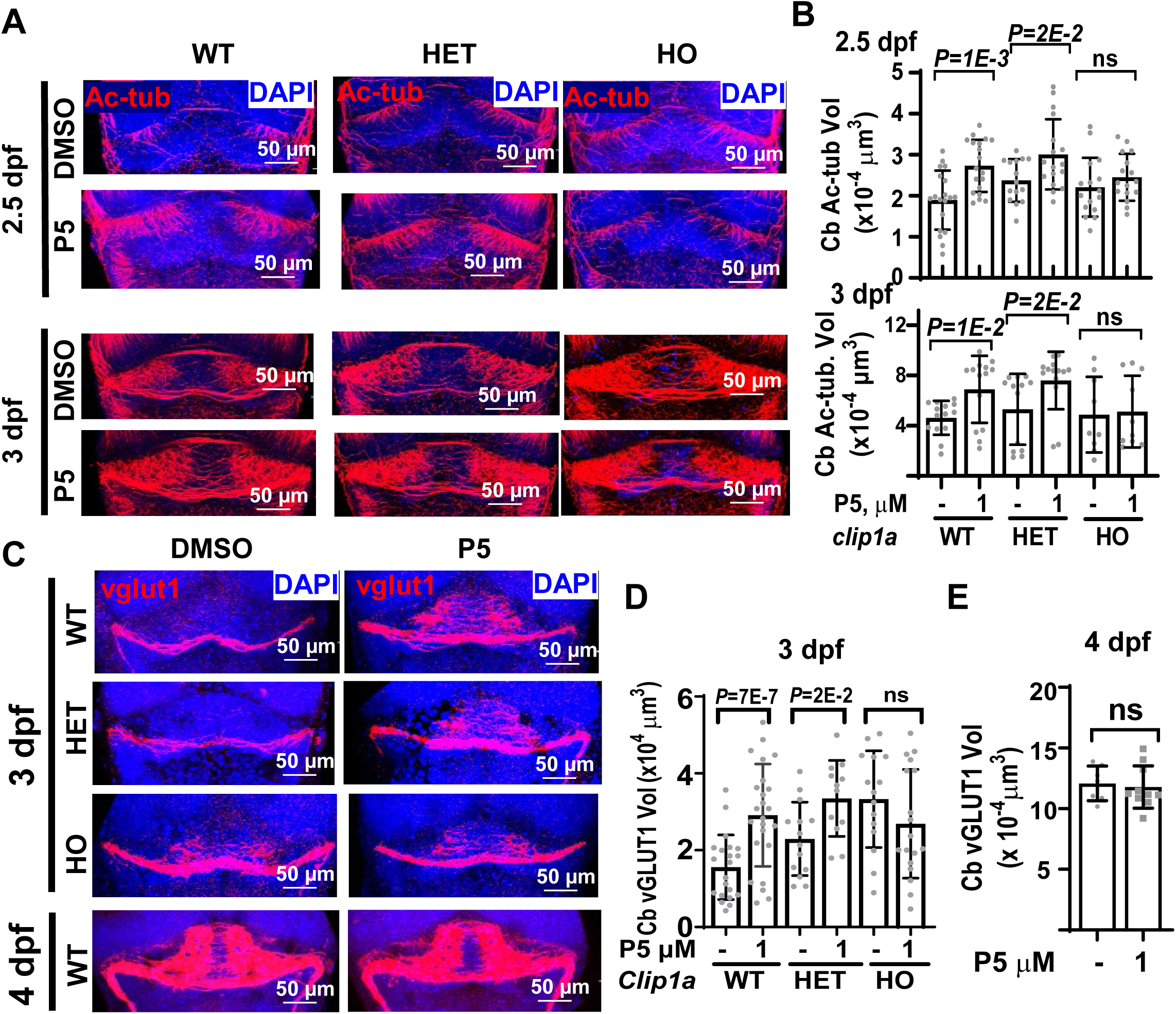
P5 promotes zebrafish cerebellum development via CLIP1. **A-B.** P5 accelerates formation of stable microtubule track in developing axons of zebrafish cerebellum via *Clip1a*; **A**. Zebrafish cerebellum immunostained with acetylated tubulin (red) at 2.5 and 3 dpf; **B**. P5 accelerates axon formation in WT and *clip1a* heterozygotes (HET), but not *clip1a* homozygous (HO) mutant embryos at 2.5 dpf F(5,96)=5.86, *p*=9*E- 5, one-way ANOVA and at 3 dpf F(5,68)=3.08, *p*=0.01, one-way ANOVA. **C-E**. P5 accelerates synaptic system formation in zebrafish cerebellum at 3 dpf; **C**. Zebrafish cerebellum immunostained with vGlut1 (red) that marks axonal synaps and DAPI (blue) at 3 dpf. **D**. P5 enhances glutamatergic axons formation in WT and *clip1a* HET cerebellum, but not in *clip1a* HO mutant cerebellum F(5,98)=6.02, *p*=6*E-5, one-way ANOVA. **E.** P5 has no effect on 4-dpf axons. Data are presented as mean±SD.

However, axons of the granule neurons at 4 dpf were indistinguishable between control and P5 treated embryos (12 ± 1 x10^-4^ µm^3^ vs 12 ± 1.7 x10^-4^ µm^3^, *p*=0.7, *t* test Fig. 9E), suggesting that CLIP1-P5 affects axon formation only during a specific time window of embryonic development.

## Discussion

### Conserved effects of P5 on neurites and axon outgrowth in different types of neurons

In this study we used four types of developing neuronal culture that are bipolar or multipolar to analyze the effect of P5. P5 increased the outgrowth of early neurites in all neurons (Fig. 1). The range of effective concentrations (0.01-1 μM) for all neuronal cell types is similar. Furthermore, both cerebellar granule neurons and hippocampal neurons respond to the same concentration of P5 (1 μM) for axon specification and outgrowth (Fig.2). This indicates that P5 exerts similar effects on neurite outgrowth, axon specification and axon outgrowth across a wide range of neuron types.

Despite similar effective concentration range of P5 for many aspects of neuron development, there is one difference. Minor neurites of hippocampal neurons respond to 0.1 μM P5 for growth in contrast to nascent axons, which respond to 1 μM P5 (Fig.2). In multipolar neurons, when future axons continue to grow, microtubules in their growth cones become more stable and F-actin density is decreased (Bradke and Dotti, 1999; Witte et al., 2008). Post-translational modifications of tubulin is not compartment-specific in minor neurites, which therefore respond to lower P5 concentration. The difference of the cytoskeleton distribution along the nascent axons and minor neurites can explain their differential response to P5.

We find that higher concentration of P5 (3 μM) is ineffective for both axon and minor neurite outgrowth (Figs. 1 and 2). Similar concentration dependence has been observed in zebrafish embryonic migration; while 0.1 and 1 μM P5 promote epiboly migration, 20 μM P5 has no effect (Hsu et al., 2006). P5 improves mouse memory performance also only at a dose window of 200 pg/mouse while being ineffective at lower and higher concentrations (Ducharme et al., 2010). Thus, P5 functions through a narrow concentration range.

### P5-CLIP1 accelerates microtubule network maturation in developing neurons

We found that P5 promotes neurite outgrowth via CLIP1, but zebrafish *Clip1a* knockout neurons have normal neurites, suggesting that CLIP1 is not necessary for neurites initiation and elongation (Fig.3). CLIP1 and CLIP2 (CLIP115) both play roles in neuronal polarization and stabilization of growth cone microtubule dynamic (Neukirchen and Bradke, 2011). CLIP2 partially replaces CLIP1. However, P5 accelerates microtubule polymerization through the activation CLIP1, but not through CLIP2 (Weng et al., 2013). CLIP1 preferentially binds tyrosinated tubulin (Bieling et al., 2008). Our data show that CLIP1 activated by P5 increases polymerization speed and penetration of dynamic microtubules into the filopodia, resulting in cytoskeleton reorganization in the growth cone and acceleration of neurite elongation.

Axon formation is associated with compartment-specific distribution of dynamic and stable pool of microtubule. The distribution of acetylated tubulin is elevated along the axonal shaft, while tyrosinated tubulin are enriched close to growth cone (Song and Brady, 2015; Nirschl et al., 2016). We detected that P5 further deepens this differential distribution of tubulin modifications. The S311D-CLIP1, which is activated by P5, confirms that dynamic/stable microtubule balance is regulated by CLIP1 activation. Thus, P5 accelerates maturation of microtubule network along nascent axon to promote axon specification.

### P5 reorganizes cytoskeleton and accelerates brain development

Our results from zebrafish cerebellum development showed that P5/CLIP1 stabilized microtubules in developing axons, accompanied with accelerated synaptic formation. This effect is observed only during a narrow period of cerebellar development, but it does not interfere with the normal developmental process. These results suggest that P5/CLIP1 can underly the beneficial role of P5 for treatment of neurodevelopmental diseases, including schizophrenia, autism spectrum disorders.

The activation of CLIP1 by P5 leads to cytoskeleton reorganization in neurons. P5 increases dynamic microtubules along newly formed axon, and promotes microtubule invasion and protrusion into the distal half of filopodia. The protrusion of microtubule into the peripheral domain of growth cones facilitates regeneration of adult DRG (Hur et al., 2011). Tubulin tyrosination is associated with increased initiation of axon regeneration (Song et al., 2015). Altogether it suggests that P5 can work along with axon regeneration processes in injured brain.

In summary, we showed that P5 through CLIP1 activation promotes early neurite outgrowth of several neuronal cell types in a concentration-dependent and neuron stage-specific manner. It accelerates microtubule dynamics and cytoskeleton reorganization, which are intrinsic for axon formation, and brain development.

## Acknowledgments

This work was funded by grants from Academia Sinica (AS NB 2396-106-0100 and AS-107-TP-L08), National Health Research Institutes (NHRI- EX107-10506SI), and Ministry of Science and Technology (MOST 107-2321-B-001-034). We would like to thank Florian Pelzer and Ching-Ju Lin for the assistance in the characterization of *clip1a* knockout fish, J.R. Henley and Han Lee for comments on zebrafish embryo dissection, and the Imaging Core (Institute of Molecular Biology Academia Sinica) for assistance in microscopy.

## References

Abdel-Hafiz L, Chao OY, Huston JP, Nikolaus S, Spieler RE, de Souza Silva MA, Mattern C (2016) Promnestic effects of intranasally applied pregnenolone in rats. Neurobiol Learn Mem 133:185–195.

Ayatollahi A, Bagheri S, Ashraf-Ganjouei A, Moradi K, Mohammadi MR, Akhondzadeh S (2020) Does Pregnenolone Adjunct to Risperidone Ameliorate Irritable Behavior in Adolescents With Autism Spectrum Disorder: A Randomized, Double-Blind, Placebo-Controlled Clinical Trial? Clin Neuropharmacol 43:139–145.

Bae YK, Kani S, Shimizu T, Tanabe K, Nojima H, Kimura Y, Higashijima S, Hibi M (2009) Anatomy of zebrafish cerebellum and screen for mutations affecting its development. Dev Biol 330:406–426.

Barbiero I, De Rosa R, Kilstrup-Nielsen C (2019) Microtubules: A Key to Understand and Correct Neuronal Defects in CDKL5 Deficiency Disorder? Int J Mol Sci 20.

Barbiero I, Peroni D, Tramarin M, Chandola C, Rusconi L, Landsberger N, Kilstrup-Nielsen C (2017) The neurosteroid pregnenolone reverts microtubule derangement induced by the loss of a functional CDKL5- IQGAP1 complex. Hum Mol Genet 26:3520–3530.

Barbiero I, Peroni D, Siniscalchi P, Rusconi L, Tramarin M, De Rosa R, Motta P, Bianchi M, Kilstrup-Nielsen C (2020) Pregnenolone and pregnenolone- methyl-ether rescue neuronal defects caused by dysfunctional CLIP170 in a neuronal model of CDKL5 Deficiency Disorder. Neuropharmacology 164:107897.

Beaudoin GM, 3rd, Lee SH, Singh D, Yuan Y, Ng YG, Reichardt LF, Arikkath J (2012) Culturing pyramidal neurons from the early postnatal mouse hippocampus and cortex. Nat Protoc 7:1741–1754.

Bieling P, Kandels-Lewis S, Telley IA, van Dijk J, Janke C, Surrey T (2008) CLIP-170 tracks growing microtubule ends by dynamically recognizing composite EB1/tubulin-binding sites. J Cell Biol 183:1223–1233.

Bradke F, Dotti CG (1999) The Role of Local Actin Instability in Axon Formation. Science 283:1931.

Cai H, Zhou X, Dougherty GG, Reddy RD, Haas GL, Montrose DM, Keshavan M, Yao JK (2018) Pregnenolone-progesterone-allopregnanolone pathway as a potential therapeutic target in first-episode antipsychotic-naive patients with schizophrenia. Psychoneuroendocrinology 90:43–51.

Calogero AM, Mazzetti S, Pezzoli G, Cappelletti G (2019) Neuronal microtubules and proteins linked to Parkinson’s disease: a relevant interaction? Biol Chem 400:1099–1112.

Dixit R, Barnett B, Lazarus JE, Tokito M, Goldman YE, Holzbaur EL (2009) Microtubule plus-end tracking by CLIP-170 requires EB1. Proc Natl Acad Sci U S A 106:492–497.

Ducharme N, Banks WA, Morley JE, Robinson SM, Niehoff ML, Mattern C, Farr SA (2010) Brain distribution and behavioral effects of progesterone and pregnenolone after intranasal or intravenous administration. Eur J Pharmacol 641:128–134.

Fontaine-Lenoir V, Chambraud B, Fellous A, David S, Duchossoy Y, Baulieu EE, Robel P (2006) Microtubule-associated protein 2 (MAP2) is a neurosteroid receptor. Proc Natl Acad Sci U S A 103:4711–4716.

Frau R, Miczan V, Traccis F, Aroni S, Pongor CI, Saba P, Serra V, Sagheddu C, Fanni S, Congiu M, Devoto P, Cheer JF, Katona I, Melis M (2019) Prenatal THC exposure produces a hyperdopaminergic phenotype rescued by pregnenolone. Nat Neurosci 22:1975–1985.

Guth L, Zhang Z, Roberts E (1994) Key role for pregnenolone in combination therapy that promotes recovery after spinal cord injury. Proc Natl Acad Sci U S A 91:12308–12312.

Hahn I, Voelzmann A, Liew Y-T, Costa-Gomes B, Prokop A (2019) The model of local axon homeostasis - explaining the role and regulation of microtubule bundles in axon maintenance and pathology. Neural Development 14:11.

Hamasaki M, Matsumura S, Satou A, Takahashi C, Oda Y, Higashiura C, Ishihama Y, Toyoshima F (2014) Pregnenolone functions in centriole cohesion during mitosis. Chem Biol 21:1707–1721.

Hsu HJ, Liang MR, Chen CT, Chung BC (2006) Pregnenolone stabilizes microtubules and promotes zebrafish embryonic cell movement. Nature 439:480–483.

Hur EM, Yang IH, Kim DH, Byun J, Saijilafu, Xu WL, Nicovich PR, Cheong R, Levchenko A, Thakor N, Zhou FQ (2011) Engineering neuronal growth cones to promote axon regeneration over inhibitory molecules. Proc Natl Acad Sci U S A 108:5057–5062.

Kahn OI, Baas PW (2016) Microtubules and Growth Cones: Motors Drive the Turn. Trends Neurosci 39:433–440.

Kani S, Bae YK, Shimizu T, Tanabe K, Satou C, Parsons MJ, Scott E, Higashijima S, Hibi M (2010) Proneural gene-linked neurogenesis in zebrafish cerebellum. Dev Biol 343:1–17.

Lansbergen G, Komarova Y, Modesti M, Wyman C, Hoogenraad CC, Goodson HV, Lemaitre RP, Drechsel DN, van Munster E, Gadella TW, Jr., Grosveld F, Galjart N, Borisy GG, Akhmanova A (2004) Conformational changes in CLIP-170 regulate its binding to microtubules and dynactin localization. J Cell Biol 166:1003–1014.

Larti F, Kahrizi K, Musante L, Hu H, Papari E, Fattahi Z, Bazazzadegan N, Liu Z, Banan M, Garshasbi M, Wienker TF, Hilger Ropers H, Galjart N, Najmabadi H (2015) A defect in the CLIP1 gene (CLIP-170) can cause autosomal recessive intellectual disability. Eur J Hum Genet 23:416.

Lasser M, Tiber J, Lowery LA (2018) The Role of the Microtubule Cytoskeleton in Neurodevelopmental Disorders. Frontiers in Cellular Neuroscience 12.

Lee HS, Komarova YA, Nadezhdina ES, Anjum R, Peloquin JG, Schober JM, Danciu O, van Haren J, Galjart N, Gygi SP, Akhmanova A, Borisy GG (2010) Phosphorylation controls autoinhibition of cytoplasmic linker protein-170. Mol Biol Cell 21:2661–2673.

Letourneau PC, Shattuck TA, Ressler AH (1987) “Pull” and “push” in neurite elongation: observations on the effects of different concentrations of cytochalasin B and taxol. Cell Motil Cytoskeleton 8:193–209.

Meijering E, Dzyubachyk O, Smal I (2012) Methods for cell and particle tracking. Methods Enzymol 504:183–200.

Mellon SH, Deschepper CF (1993) Neurosteroid biosynthesis: genes for adrenal steroidogenic enzymes are expressed in the brain. Brain Res 629:283–292.

Nakano A, Kato H, Watanabe T, Min KD, Yamazaki S, Asano Y, Seguchi O, Higo S, Shintani Y, Asanuma H, Asakura M, Minamino T, Kaibuchi K, Mochizuki N, Kitakaze M, Takashima S (2010) AMPK controls the speed of microtubule polymerization and directional cell migration through CLIP-170 phosphorylation. Nat Cell Biol 12:583–590.

Naylor JC, Kilts JD, Hulette CM, Steffens DC, Blazer DG, Ervin JF, Strauss JL, Allen TB, Massing MW, Payne VM, Youssef NA, Shampine LJ, Marx CE (2010) Allopregnanolone levels are reduced in temporal cortex in patients with Alzheimer’s disease compared to cognitively intact control subjects. Biochim Biophys Acta 1801:951–959.

Naylor JC, Kilts JD, Shampine LJ, Parke GJ, Wagner HR, Szabo ST, Smith KD, Allen TB, Telford-Marx EG, Dunn CE, Cuffe BT, O’Loughlin SH, Marx CE (2020) Effect of Pregnenolone vs Placebo on Self-reported Chronic Low Back Pain Among US Military Veterans: A Randomized Clinical Trial. JAMA network open 3:e200287.

Neukirchen D, Bradke F (2011) Cytoplasmic linker proteins regulate neuronal polarization through microtubule and growth cone dynamics. J Neurosci 31:1528–1538.

Nirschl JJ, Magiera MM, Lazarus JE, Janke C, Holzbaur EL (2016) alpha- Tubulin Tyrosination and CLIP-170 Phosphorylation Regulate the Initiation of Dynein-Driven Transport in Neurons. Cell Rep 14:2637–2652.

Orefice N, Carotenuto A, Mangone G, Bues B, Rehm R, Cerillo I, Sacca F, Calignano A, Orefice G (2016) Assessment of neuroactive steroids in cerebrospinal fluid comparing acute relapse and stable disease in relapsing-remitting multiple sclerosis. J Steroid Biochem Mol Biol 159:1–7.

Perez F, Diamantopoulos GS, Stalder R, Kreis TE (1999) CLIP-170 highlights growing microtubule ends in vivo. Cell 96:517–527.

Ritsner MS, Bawakny H, Kreinin A (2014) Pregnenolone treatment reduces severity of negative symptoms in recent-onset schizophrenia: an 8-week, double-blind, randomized add-on two-center trial. Psychiatry Clin Neurosci 68:432–440.

Shin YY, Kang EJ, Jeong JS, Kim MJ, Jung EM, Jeung EB, An BS (2019) Pregnenolone as a potential candidate for hormone therapy for female reproductive disorders targeting ERbeta. Mol Reprod Dev 86:109–117.

Song W, Cho Y, Watt D, Cavalli V (2015) Tubulin-tyrosine Ligase (TTL)- mediated Increase in Tyrosinated alpha-Tubulin in Injured Axons Is Required for Retrograde Injury Signaling and Axon Regeneration. J Biol Chem 290:14765–14775.

Song Y, Brady ST (2015) Post-translational modifications of tubulin: pathways to functional diversity of microtubules. Trends in Cell Biology 25:125–136.

Tahirovic S, Bradke F (2009) Neuronal polarity. Cold Spring Harb Perspect Biol 1:a001644.

Tomaselli G, Vallee M (2019) Stress and drug abuse-related disorders: The promising therapeutic value of neurosteroids focus on pregnenolone- progesterone-allopregnanolone pathway. Front Neuroendocrinol 55:100789.

Vallée M (2016) Neurosteroids and potential therapeutics: Focus on pregnenolone. The Journal of Steroid Biochemistry and Molecular Biology 160:78–87.

Vallee M et al. (2014) Pregnenolone can protect the brain from cannabis intoxication. Science 343:94–98.

Weng JH, Chung BC (2016) Nongenomic actions of neurosteroid pregnenolone and its metabolites. Steroids 111:54–59.

Weng JH, Liang MR, Chen CH, Tong SK, Huang TC, Lee SP, Chen YR, Chen CT, Chung BC (2013) Pregnenolone activates CLIP-170 to promote microtubule growth and cell migration. Nat Chem Biol 9:636–642.

Westerfield M (2000) The zebrafish book. A guide for the laboratory use of zebrafish (Danio rerio). 4th ed, Univ of Oregon Press, Eugene.

Wilkins A, Majed H, Layfield R, Compston A, Chandran S (2003) Oligodendrocytes Promote Neuronal Survival and Axonal Length by Distinct Intracellular Mechanisms: A Novel Role for Oligodendrocyte- Derived Glial Cell Line-Derived Neurotrophic Factor. The Journal of Neuroscience 23:4967–4974.

Witte H, Neukirchen D, Bradke F (2008) Microtubule stabilization specifies initial neuronal polarization. J Cell Biol 180:619–632.

Yamamoto H, Demura T, Morita M, Banker GA, Tanii T, Nakamura S (2012) Differential neurite outgrowth is required for axon specification by cultured hippocampal neurons. J Neurochem 123:904–910.

Yang X, Li H, Liu XS, Deng A, Liu X (2009) Cdc2-mediated phosphorylation of CLIP-170 is essential for its inhibition of centrosome reduplication. J Biol Chem 284:28775–28782.

Young J, Corpéchot C, Perché F, Eychenne B, Haug M, Baulieu EE, Robel P (1996) Neurosteroids in the mouse brain: behavioral and pharmacological effects of a 3 beta-hydroxysteroid dehydrogenase inhibitor. Steroids 61:144–149.

Zhang F, Su B, Wang C, Siedlak SL, Mondragon-Rodriguez S, Lee H-G, Wang X, Perry G, Zhu X (2015) Posttranslational modifications of α-tubulin in alzheimer disease. Transl Neurodegener 4:9–9.

Zwain IH, Yen SS (1999) Neurosteroidogenesis in astrocytes, oligodendrocytes, and neurons of cerebral cortex of rat brain. Endocrinology 140:3843–3852.

